# Simple sequence repeats power *Staphylococcus aureus* adaptation

**DOI:** 10.1101/2025.07.08.663602

**Authors:** Romain Guérillot, Ashleigh Hayes, Abderrahman Hachani, Stefano G. Giulieri, Liam Donovan, Louise M. Judd, Liam K. Sharkey, Anders Gonçalves da Silva, Torsten Seemann, Timothy P. Stinear, Benjamin P. Howden

## Abstract

Understanding *Staphylococcus aureus* adaptation is foundational for addressing the challenges posed by this pathogen. Using evolutionary convergence analysis on 7099 *S. aureus* genomes we uncovered frequent, reversible loss-of-function (LoF) mutations caused by simple sequence repeats (SSRs). Functional assays showed SSRs are potent, length-dependent switches that activate and deactivate genes, a mechanism exemplified by frequent LoF mutations in *mutL* (DNA mismatch repair) that causes a hypermutator phenotype. Analysis of over 600 episodes of human *S. aureus* colonisation and infection confirmed laboratory findings and showed that *mutL* SSR hypermutability behaves like a phase-variable evolutionary capacitor that unlocks subsequent adaptive SSR variations. We also experimentally demonstrate that SSRs facilitate foreign DNA uptake (*hsdR* restriction endonuclease) and lead to gentamicin-resistant small colony variants that are known to cause persistent infections *(menF* menadione biosynthesis). Furthermore, the phenomenon is widespread, with 27 of 29 diverse human microbiome samples harbouring *S. aureus* with distinct SSR subpopulations, indicative of niche-specific adaptations. Thus, SSRs are potent reservoirs of previously unrecognized *S. aureus* genetic heterogeneity that confer rapid adaptive capacity, including in antigens targeted by past vaccine trials. A deeper understanding of SSR evolutionary trajectories has the potential to improve treatments and to anticipate *S. aureus* responses to new therapies.

## Introduction

Convergent evolution is a fascinating feature of adaptation, where independently evolving lineages develop identical solutions to cope with similar selective pressures ^1,2^. Identifying and characterizing genetic convergence in microbial pathogens is a powerful technique for exposing the strategies microorganisms depend upon to survive and proliferate ^3–6^. Mutations conferring antimicrobial resistance are prototypical examples of convergent evolution (also called homoplasy) ^7–10^. Identifying instances of convergent evolution has also revealed important host adaptations mechanisms during infection ^11–13^ and immune-evasion associated with widespread transmission ^14,15^.

*Staphylococcus aureus* is an opportunistic bacterial pathogen of global importance, causing high morbidity and mortality ^16^. The bacterium is notorious for rapidly adapting to changing environments and causing a spectrum of infections, ranging from mild skin and soft tissue infections to life-threatening bloodstream and cardiac infections ^17^. The adaptability of *S. aureus* stems from its large repertoire of genetic and phenotypic variations. The *S. aureus* genome encodes a diverse repertoire of virulence determinants, including toxins, adhesins, and immune evasion molecules whose expression is controlled by regulatory systems ^18,19^. *S. aureus* can rapidly change to different phenotypic states ^20–22^. These changes in infecting population cause hetero-resistance to antibiotics and other pathoadaptations that compromise patient treatments ^23,24^. For example, subpopulations of *S. aureus* can develop a persister phenotype by halting bacterial growth and survive immune or antibiotic clearance. When isolating *S. aureus* from infections, a similar but directly observable phenotype is the Small Colony Variant (SCV). SCVs are slow growing *S. aureus* subpopulations that usually show decreased haemolytic activity, increased antibiotic resistance, and persistence within cells. SCVs and other phenotypic switching during infection have serious clinical implications as these changes are linked with treatment failures and relapsing infections. However, the molecular bases of these phenomena remain poorly understood ^25,23^.

Convergent resistance mutations resulting from antibiotic selection are frequently observed in bacterial pathogens ^7,8,12,26^. With advances in genomics and sequencing technologies, it is now possible to compare thousands of *S. aureus* genomes to detect genetic convergence with unprecedented sensitivity and uncover adaptations resulting from less stringent selective pressures. Identifying the most convergent mutations and characterising their impact during infections can provide a better understanding of what is driving pathogenesis and ultimately help develop new strategies to fight infections.

Here we performed a large-scale evolutionary convergence analysis across 7,099 publicly available genomes and reveal the important role played by Simple Sequence Repeats (SSRs) mediated frameshifts in *S. aureus* adaptability. By pinpointing the most frequently recurring signals of convergent evolution, we identify conserved SSR loci as a central molecular mechanism driving a large spectrum of phenotypically diverse subpopulations of *S. aureus.* We found niche-specific subpopulations with different patterns of SSR variation in human microbiome-derived metagenomes, variation that was missed by sequencing single *S. aureus* isolates. Using experimental evolution, *ex vivo* infection model and clinical *S. aureus* samples, we demonstrate that these SSR driven subpopulations can adapt faster, resist antibiotics and swiftly switch normal interactions with the host.

## Results

### Convergent Mutation Adaptive Score identifies adaptation signatures in *S. aureus*

To identify the genetic loci with the highest potential for adaptive evolution, we quantified all occurrences of convergent mutations across a global phylogeny of 7099 *S. aureus* genomes (Fig. 1a,b). For the 573,721 unique mutations detected in this population, we counted the minimum number of independent, monophyletic acquisition events for each mutation using parsimony analysis on the core-genome SNP tree representing the evolutionary history of all *S. aureus* strains (Fig. 1c). To pinpoint the most likely adaptive mutations, we developed a *Convergent Mutation Adaptive Score* (CMAS) that standardised the number of independent, non-synonymous mutation acquisitions with all mutation acquisitions for every gene. For each non-synonymous mutation, CMAS measures how many standard deviations its independent-acquisition count exceeds the mean acquisition count across all sites in the same gene. This standardisation of acquisition counts at the gene-level corrects for linkage disequilibrium due to recombination—where blocks of mutations co-transfer and would otherwise appear as convergent mutations—and prevents these linked acquisition events from inflating convergent adaptive signals. It also accounts for inherent variability in mutation rates across genes arising from gene length, divergence time, and functional constraints, ensuring that only mutations with acquisition counts truly exceeding their expected variance are identified (see methods). CMAS values are then evaluated against their empirical genome-wide distribution—without relying on any parametric model. A non-parametric 99% prediction interval is then used to identify the rare mutated loci whose exceptionally high recurrence signals are indicative of positive selection. CMAS enabled the identification of 3597 highly convergent mutations with a high probability of reflecting an adaptive evolutionary response (CMAS > 99% upper boundary prediction interval, Fig. 1d, Supp. Table 1). Mutations with the highest CMAS corresponded to recurrent adaptive mutations conferring resistance to fluoroquinolone (*parC*) and rifampicin (*rpoB*) antibiotics (Fig. 1d). Among these highly convergent mutations, we found that at least 45 mutations in 14 genes confer previously described adaptative phenotypes: resistance to 8 different antibiotic families (10 genes), resistance to the biocide triclosan (1 gene), virulence (1 gene), biofilm formation (1 gene), host-interaction (1 gene; Fig. 1d and e; Supp. Table 2). These mutations arose with high frequency across the *S. aureus* population (Fig. 1c). Importantly, each point mutations with the highest CMAS within these 14 genes pinpoint well described adaptive mutations, underscoring the robustness of our mining approach for adaptation signatures (Fig. 1e; Supp. Fig. 1). These results demonstrate that CMAS identifies highly convergent point mutations causing adaptive phenotypes from a large collection of *S. aureus* genome without requiring information other than DNA sequences.

**Figure 1.**
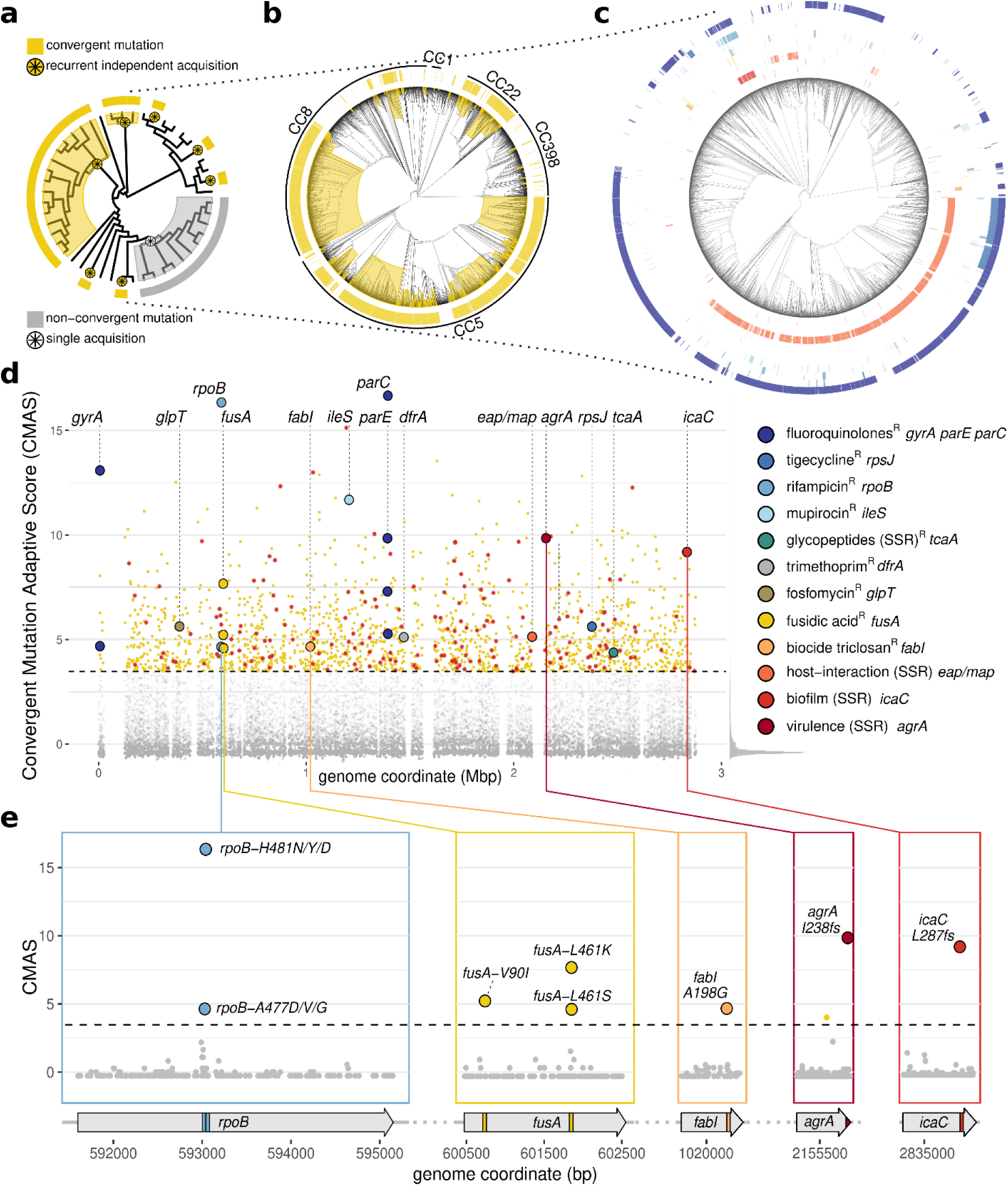
Convergent Mutation Adaptive Score analysis (CMAS). **a**. Circular phylogenetic tree illustrating a convergent mutation (yellow) that has been recurrently acquired 7 times during evolution (yellow circled asterisk) and a non-convergent mutation (grey) with a single acquisition (black circled asterisk). **b**. Circular phylogenetic tree of 7099 S. aureus isolates inferred from core-genome SNP. Branch length Major Clonal Complexes (CC) are represented together with an example of a highly convergent mutation (yellow). **c**. Tree displaying mutation presence/absence heatmap of highly recurrent known adaptive mutations re-identified with CMAS analysis; mutation colour legend matches panel d legend. The branch lengths of the trees have been unscaled for visualization **d**. Genome-wide CMAS plot. Mutations with significantly high CMAS (>99% prediction interval, dashed line) are represented by yellow dots (non-synonymous mutations) and red asterisk (Simple Sequence Repeat [SSR]-mediated mutations). Mutations previously known to confer adaptive phenotypes prior to this study are indicated with coloured circles. corresponding phenotypes indicated in the legend. Empirical distribution of CMAS is represented on a vertical grey histogram on the right side of the CMAS plot: most mutations form a low-score peak, while the dashed prediction interval 99% cutoff slices off the extreme right tail capturing the small subset of mutations with exceptionally high convergence. **e.** Gene-wide CMAS plots. Point mutations known to confer adaptation prior to this study consistently exhibit significantly high CMAS score (>99% CMAS prediction interval, dashed line). The mutation colour legend is consistent with the legend in panel d.

### Highly convergent mutations arise frequently within Simple Sequence Repeats (SSRs) loci causing frameshift mutations

By investigating the type of mutations associated with a significant CMAS, we found that Loss-of-Function (LoF; *i.e.* frameshifts mutations, premature stop codons, loss of start codons) were significantly overrepresented, compared to other types of mutations (n=500, p≤0.001, OR = 1.5, one-sided Fisher’s exact test; Supp. Table 1). Interestingly, recurrent LoF mutations have been previously observed across different collections of invasive *S. aureus* isolates and from within-host evolution studies ^12,27–29^. This observation prompted us to investigate the genetic basis for these events. We discovered that the most significant enrichment corresponded to frameshift mutations arising within DNA sequences and consisting of repeated units of one (n=175), two (n=2) or four nucleotides (n=2) (p≤ 0.001, OR=3.6, one-sided Fisher’s exact test; Fig. 2a). These repeats, known as Simple Sequence Repeats (SSRs), are prone to insertion/deletion (indel) mutations by DNA polymerase slippage during DNA replication (Fig. 2b) ^30^. We noted that SSR-mediated frameshift mutations represented 76% (379 of 500) of all highly convergent LoF mutations. These highly convergent SSR frameshifts recurred across 186 different SSR loci and 171 different genes (Supp. Table 3). Three previously reported adaptive SSR-mediated frameshift mutations in the genes *agrA*, *icaC* and *eap/map* illustrate the phenomenon and underline the validity of CMAS (Fig. 1c, d and e) ^31,32,24,33,13^. We also observed that as SSR length increased, so did the frequency of adaptive mutations. SSRs of length >5 nucleotides had significantly higher CMAS and mutation acquisition than non-SSR indel and SNP mutations (p≤0.0001, two-sided Wilcoxon test), with both metrics increasing with the length of the SSR (Fig. 2c and d).

**Figure 2.**
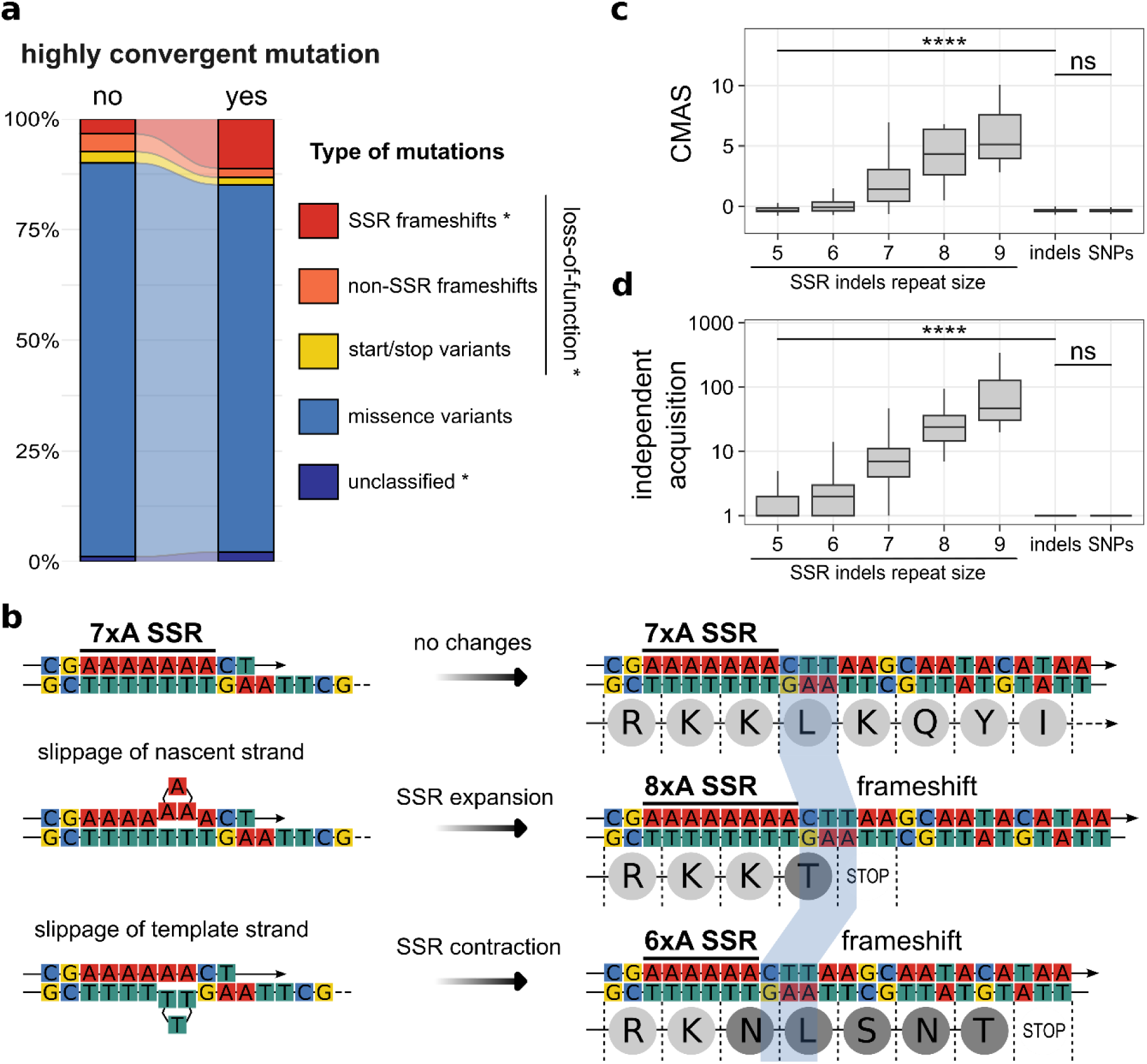
SSR-frameshifts cause highly convergent mutations in S. aureus. **a.** Stacked bar chart representing the significant enrichment of LoF mutations mediated by SSR-frameshifts among highly convergent mutations (Supp. Table 1). * indicate that the mutation categories loss-of-function, SSR-frameshift and unclassified are significantly enriched in highly convergent mutations compared to non-highly convergent mutations (p<0.001 one-sided Fisher’s exact test). **b.** Replication slippage induces SSR expansion or contraction in nucleotide repeats. These cause frameshift and premature stop codon during the translation of coding sequences – leading to truncated proteins and LoFs. **c.** CMAS and **d.** number of independent acquisition of highly convergent SSR-indels increase with the size of SSR repeats and are significantly higher than non SSR-mediated indels and SNPs (p≤0.0001, two-sided Wilcoxon test).

### SSRs cause a high rate of reversible frameshift mutations

To experimentally assess the impact of SSR length on mutation rate, we constructed *S. aureus* strains with chromosomal encoded reporters of SSR contractions and expansions. We introduced seven artificial SSRs, consisting of five to nine adenosine nucleotide repeats ((A)_5_ to (A)_9_) into the *ermC* gene, that confers resistance to macrolide antibiotics ^34^. Each of these seven SSRs disrupted the functionality of the *ermC* allele by introducing a +1 or -1 frameshift mutation through the addition or deletion of an adenine nucleotide. Consequently, the contraction or expansion of an SSR by one nucleotide restores the functionality of *ermC*, resulting in an antibiotic resistance phenotype (Fig. 3a). We measured the frequency and mutation rate of SSR expansion and contraction for SSR repeats by plating the strains on media with and without azithromycin (Fig. 3b). We found that all SSR expansions and contractions arose at frequencies at or above rifampicin resistance mutation frequency. Rifampicin is an antibiotic for which resistance can be selected at a high rate by single-step selection and that does not involve SSRs (Supp. Fig. 2) ^35,36^. As observed from our population genomic investigation, mutation frequencies increased significantly with SSR length. For every repeat length tested, SSRs were more prone to expansion than contraction (Fig. 3c).

**Figure 3.**
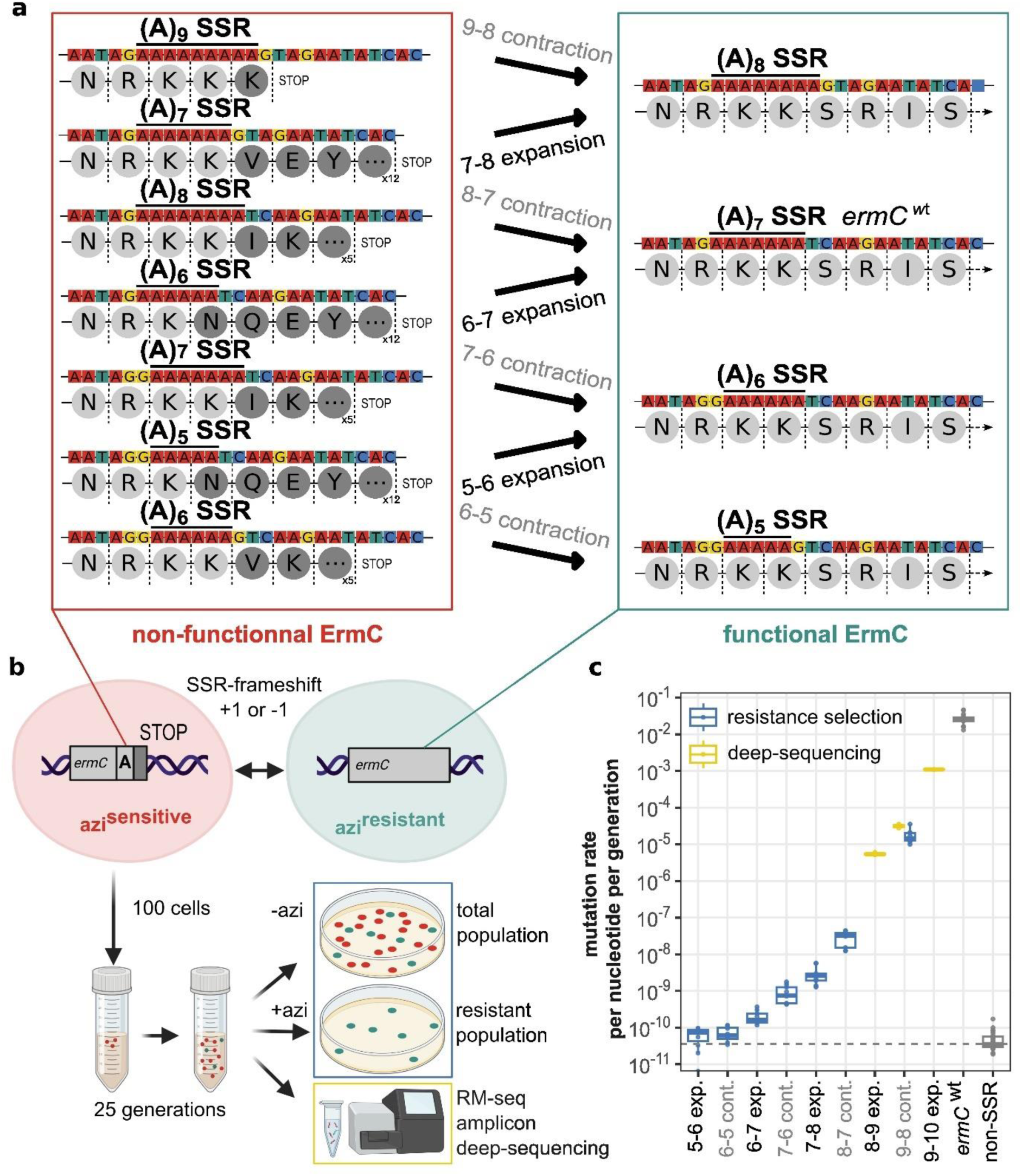
Azithromycin resistance reporter assay to measure the rate of SSR-mediated frameshifts. **a.** Sequences of synthetic reporter of SSR-mediated contraction and expansion. Left: The seven reporters constructed introduced a +1 or –1 frameshift in a functional ermC resistance gene for SSRs of repeat size 5 to 9, causing changes in protein sequence (dark grey amino acid) and premature stop codons. Right: Contraction or expansion of the SSR by 1 nucleotide reconstitute a functional ermC gene. **b.** Bacterial cells with SSR-frameshifted ermC allele are sensitive to azithromicin antibiotic (azi^sensitive^) and –1 or +1 SSR-frameshift render cells resistant to azihromycin (azi^resistant^). After culture in broth of a low inoculum of azi^S^ reporter cells, cultures are plated onto selective (+azi) and non-selective (-azi) agar. Colonies are enumerated to count resistant subpopulation and total population and to calculate frequencies and rates of SSR-frameshifts of the different reporters (blue frame). Alternatively, after broth culture genomic DNA is extracted and RM-seq amplicon deep-sequencing method is used to directly count the abundance of wild type and frameshifted SSRs (yellow frame) ^36^. **c.** Mutation rates per nucleotide and per generation of the 7-nucleotide SSR expansions (exp.) and contractions (cont.) assessed using antibiotic resistance selection (n=9) and/or amplicon deep-sequencing (n=2). Non-SSR (i.e. basal) mutation rate per nucleotide per base of S. aureus was inferred from the frequency of rifampicin resistance (n=23). A reporter strain with ermC wild type gene was used as a control (n=19).

Given the high mutation frequency for the longest SSRs (SSR contraction *ermC-*(A)_9→8_; median mutation frequency: 2.5×10^-3^), we were able to measure mutation frequencies by directly sequencing the genomic DNA of *S. aureus* cultures without selection. We applied RM-seq, an amplicon deep-sequencing method that uses molecular barcoding enabling the accurate quantification of rare allelic variants in heterogeneous populations ^36^. Using RM-seq, we measured the (A)_8→9_ and (A)_9→10_ mutation expansion frequencies by azithromycin resistance selection (median: 7.8×10^-4^ and 1.6×10^-1^, respectively) and confirmed the (A)_9→8_ contraction frequency (median: 5.1×10^-3^; Supp. Fig. 2).

For contraction and expansion of SSR repeats size >6x, the mutation rates calculated per generation and per base were significantly higher than non-SSR mutation rates (p≤0.05, one-sided Wilcoxon test; see method), with an increase up to 2.2×10^7^ times for the (A)_9→10_ expansion (Supp. Fig. 2). These results show the capacity of SSR loci to cause reversible LoF mutations at high rates and therefore rapidly switch genes “on” or “off” in *S. aureus* (i.e. phase variation).

### SSR frameshift mutations in *mutL* potentiate mutation rate and adaptability

As SSRs bearing the longer repeats size mutate at very high frequency, we interrogated their impact. Among the three SSR-frameshifts carrying the longest repeat ((N)_9_ nucleotides), the recurrent *mutL-N343fs* in the gene encoding a DNA mismatch repair protein *MutL*, had the highest CMAS (Supp. Table 3). Alteration of *mutL and mutS* has been previously shown to be associated with a hypermutator phenotype in *S. aureus* ^37^. To investigate if the SSR *mutL* frameshifts increase *S. aureus* mutation frequency, we introduced the (A)_9→8_ SSR mutation in wild type *S. aureus*, to recreate the *mutL-N343fs*. As predicted, the *mutL* SSR-frameshift caused a hypermutator phenotype, characterised by a significantly increased mutation frequency (>36-fold; Figure 4a; p≤0.0001, two-sided Wilcoxon test). To assess the impact of *mutL-N343fs* on the emergence of resistance in *S. aureus*, we repeatedly exposed this mutant and its wild type counterpart to sub-inhibitory concentrations of last-line antibiotics vancomycin and daptomycin. Whilst the vancomycin minimum inhibitory concentration (MIC) in wild type *S. aureus* increased to 4 mg/L after one passage and then remained stable, the vancomycin MIC reached 16 mg/L in the *mutL* mutant after five serial passages, in three independent experiments. Similarly, the daptomycin MIC increased to 16 mg/L in the wild type strain and increased to between 64 to 128 mg/L in the *mutL-N343fs* mutant. These results demonstrate that naturally recurring SSR-mediated frameshifts in *mutL* confer enhanced ability to rapidly adapt and become resistant to critical antibiotics commonly used to treat MRSA infections (Figure 4b).

**Figure 4.**
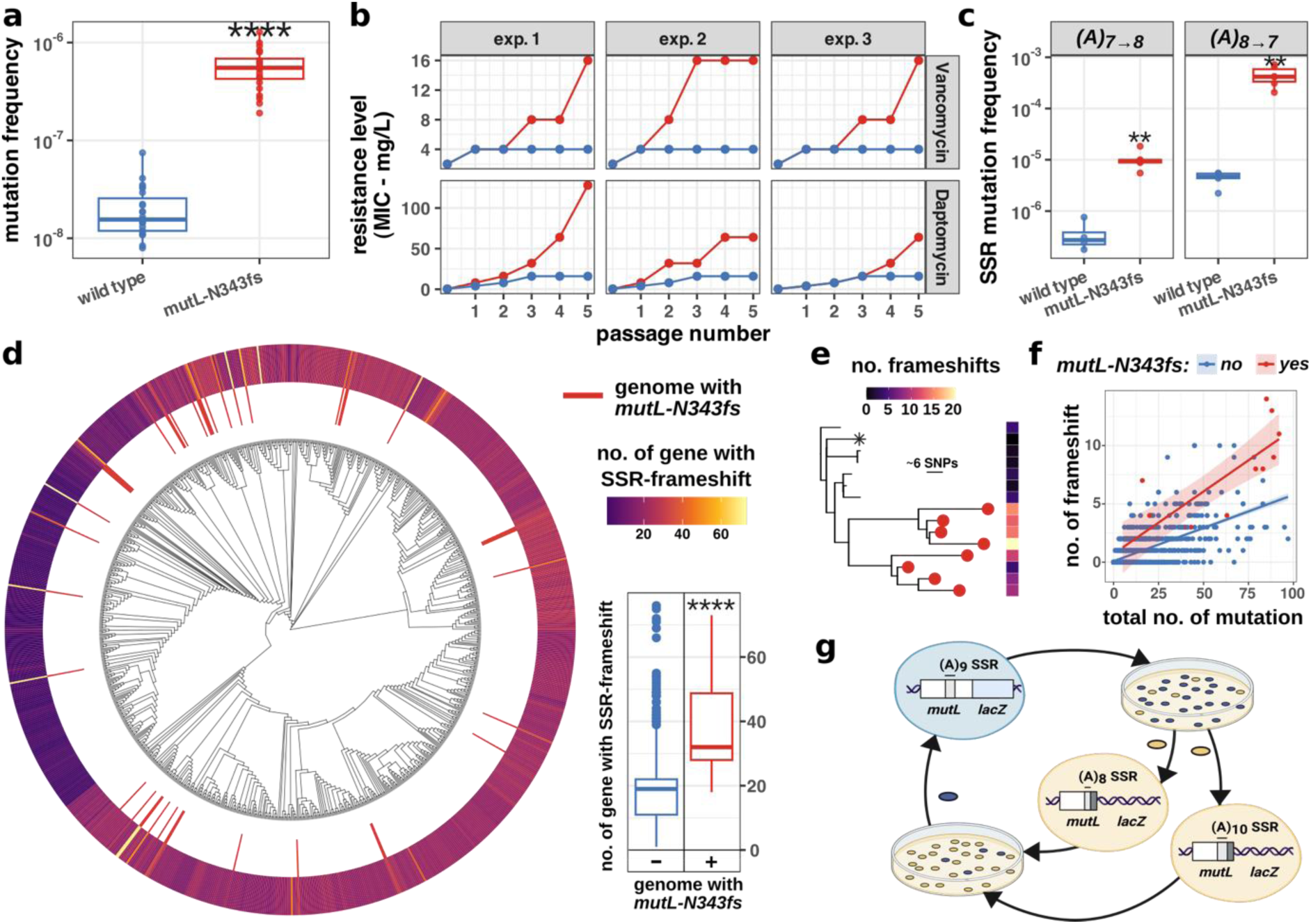
SSR-mediated frameshift in mutL increases S. aureus adaptability. **a.** Mutation frequency measured by rifampicin spontaneous resistance of wild type versus mutL-N343fs. **b.** Increase in resistance (MIC) to the antibiotics vancomycin (top) and daptomycin (bottom) of 3 independent experiments of serial exposure with a sub-inhibitory concentration of antibiotics for the mutL-N343fs strain (red line) versus wild type (blue line). Mutation detected by whole genome sequencing of isolate from the last exposure are listed in Supp. Table 4. **c.** Effect of the mutL-N343fs on frequencies of SSR expansion and contraction. SSR mutation frequencies of the ermC reporter (A)_7→8_ and (A)_8→7_ in the NRS384 wild type strain (blue) and NRS384-mutL-N343fs background (red). **d.** Sequenced S. aureus strains with an SSR-mediated frameshift in the mutL gene have significantly more SSR-frameshifts in their genomes (**** p≤0.0001, two-sided Wilcoxon test). **e.** Reconstruction of within-host evolutionary trajectories of one case of S. aureus airway infection in cystic fibrosis. The maximum likelihood tree was inferred from a core genome alignment with the internal reference isolate (asterisk). Red tips indicate isolates with mutL-N343fs SSR mutation. The heatmap shows the number of within-host acquired frameshifts. **f.** Correlation between total number of within-host acquired mutations and number of within-host acquired frameshifts stratified by presence of a within-host acquired mutL-N343fs. The lines show the fitted linear regression (mean predicted value and 95% confidence interval). **g.** Construction of a mutL-lacZ gene fusion reporter strain demonstrates the spontaneous SSR-mediated frameshift in mutL, and its reversion (i.e. phase-variation). Upon plating of a culture of a single blue colony (X-gal supplemented plates) of the fusion reporter strain mutL-lacZ-N343-(A)_9_, display a mixed population of blue and white colonies. White colonies correspond to SSR mediated frameshift in mutL-lacZ gene fusion (mutL-lacZ-N343fs-(A)_8_ or mutL-lacZ-N343fs-(A)_10_). Plating sub-cultures of both types of white colonies generate subpopulations of blue colonies that correspond to frameshift reversion to wild type sequence of the mutL-lacZ fusion (mutL-lacZ-N343-(A)_9_).

### *mutL* SSR frameshift acts as an evolutionary capacitor and emerges during *S. aureus* infection and colonisation

Whole genome sequencing of isolates arising from the last daptomycin passage (Figure 4b) showed a high prevalence of other SSR frameshift mutations in the *mutL-N343fs* background, as compared to wild type. Across three *S. aureus* isolates from independent daptomycin exposure experiments, we identified eight SSR-mediated frameshifts among 32 mutations (Supp. Table 4). In comparison, no SSR frameshifts were observed in the same number of wild type isolates sequenced after passaging. This observation prompted us to assess the effect of the *mutL* SSR-frameshift on the frequencies of subsequent SSR mutations. We compared the SSR expansion and contraction frequencies in *mutL-N343fs* versus *mutL wild-type* strain using our *ermC* reporters (*ermC-(A)_7→8_* and *(A)_8→7_*). A significant increase in the frequencies of SSR mutations (p≤0.0001, two-sided Wilcoxon test) was detected in the presence of the *mutL* frameshift, with up to an 89-fold increase in SSR-frameshifts frequencies for the reporter *ermC-(A)_8→7_* (Fig. 4c). We then examined if the presence of *mutL-N343fs* in *S. aureus* genomes was associated with an overall increase in SSR-frameshifts. There was a significant correlation between the presence of *mutL-N343fs* and the total number of SSR-frameshifts (Fig. 4d; r=0.2; p<0.001; point-biserial correlation). Furthermore, the number of SSR-frameshifts in strains carrying the *mutL-N343fs* was also significantly higher than in strains where this SSR mutation was absent (Fig. 4d; p≤0.0001, two-sided Wilcoxon test).

To further investigate the clinical relevance of the SSR-mediated *mutL-N343fs*, we performed a within-host evolution analysis of a collection of 2,189 genomes obtained from 626 episodes of *S. aureus* nasal colonisations or airway infections of cystic fibrosis patients. We detected the emergence of this mutation in seven episodes of cystic fibrosis (p≤0.0001; Poisson regression) and three episodes of nasal colonisation (p≤0.0001) (Supp. Table 5). In at least one episode, the emergence of *mutL-N343fs* sub-lineage led to a burst of up to 20 other frameshift mutations in the descendants (including in genes relevant for virulence [*sigB*] and antibiotic resistance [*dfrA*]) (Fig. 4e). To investigate whether the enrichment of frameshifts in the *mutL-N343fs* strain was also observed in the clinical context, we considered the correlation between the total number of mutations and number of frameshifts in *mutL-N343fs^+^* and *mutL-N343fs^-^* isolates. We found that the *mutL-N343fs* was associated with an almost 3-fold increase in frameshifts (Fig. 4f; β=2.97; p≤0.0001, two-sided t-test; Supp. Table 5).

These results indicate that the *mutL* SSR-mediated frameshift acts as master switch that amplifies mutation rate and triggers genome-wide SSR-mediated adaptation during colonisation or infection. By reducing the normal repair of SSR frameshifts, *mutL-N343fs* thus plays the role of an evolutionary capacitor that can unlock spectrum of potentially adaptive SSR variations ^38^.

### Hypermutator subpopulations are continually generated *via mutL* SSR fluctuations

We next assessed the proportion of *S. aureus* cells in culture harbouring SSR-frameshifted *mutL* using RM-seq deep-sequencing. We detected *mutL-N343fs* at a frequency of 1.3×10^-1^, a frequency similar to that observed for the *erm*C*-(A)_9_* SSR reporter having same SSR size (Supp. Fig. 2a, Fig. 3c). We next developed a non-selective phenotypic screen exploiting a chromosomal in-frame fusion of the wild type *mutL* with the *lacZ* reporter gene (encoding β-galactosidase) to assess the *mutL* SSR fluctuation. Growth of the *S. aureus mutL-lacZ* strain on media supplemented with X-gal (a substrate turning blue upon cleavage by β-galactosidase) showed a subpopulation of white colonies amidst blue colonies (Fig. 4g). Sequencing of *mutL*-SSR of 10 white colonies revealed that all colonies corresponded to *mutL-N343fs* mutants, with two presenting a SSR expansion (A)_9→10_ and eight corresponding to a SSR contraction (A)_9→8_. We then grew *mutL-lacZ-N343fs-(A)_8_*and *mutL-lacZ-N343fs-(A)_10_* and observed subpopulations of blue colonies among white colonies, demonstrating reversion of the mutation to the initial in-frame *mutL-lacZ* fusion (Fig. 4g). These results show that, *via mutL* SSR, *S. aureus* readily generates subpopulation of hypermutator cells that revert to wildtype. To our knowledge, this is the first evidence of a phase-variation mechanism controlling mutation rate in *S. aureus*.

### SSR-frameshift modulates horizontal gene transfer capacity

Recurrent frameshifts in the conserved *hsdR* gene, encoding the endonuclease of Type I Restriction-Modification System (T1-RMS) ^39^, were also amongst the highly convergent SSR-frameshift identified in the global collection of *S. aureus* genomes. T1-RMS prevent the acquisition of foreign DNA by transformation, conjugation and transduction in *S. aureus* ^40^and are thought to have evolved as a bacterial defence mechanism against lytic phage infections. However, acquisition of foreign DNA (*i.e.* horizontal gene transfer) is also a major driving force for bacterial evolution and often promotes adaptation ^41^, as exemplified through the transfer of the *mecA* gene, conferring *S. aureus* with methicillin resistance ^42^. The T1-RMS in *S. aureus* protects the bacterium from foreign DNA by recognising and cleaving DNA sequences bearing non-cognate methylation patterns. Thus, the *hsdR-M900fs* identified by CMAS (Supp. Table 1, 3) is predicted to increase the ability of *S. aureus* to acquire foreign DNA. To test this, we introduced the *hsdR-M900fs* mutation in a wild type background. We observed an 822-fold increase in DNA transformation efficiency in this mutant compared to the wild type. Complementation of the mutant with a wild type copy of *hsdR* restored transformation efficiency to wild type levels, demonstrating that the SSR mutation affecting *hsdR* led to increased foreign DNA uptake (Fig. 5). Plasmids pre-methylated by T1-RMS methylase to mimic non-foreign DNA were transformed at a high frequency, equivalent to that of *hsdR-M900fs* in absence of pre-methylation, in the wild type, *hsdR-M900fs* mutant and complemented mutant. This confirms that *hsdR*-SSR is specifically controlling the horizontal transfer of non-self-methylated foreign DNA. This finding illustrates how *S. aureus* has evolved a SSR in *hsdR* that generates a subpopulation of cells that can efficiently acquire potentially beneficial exogeneous DNA, while keeping most of the population protected from potential deleterious effect of foreign DNA (e.g. from lytic phages).

**Figure 5.**
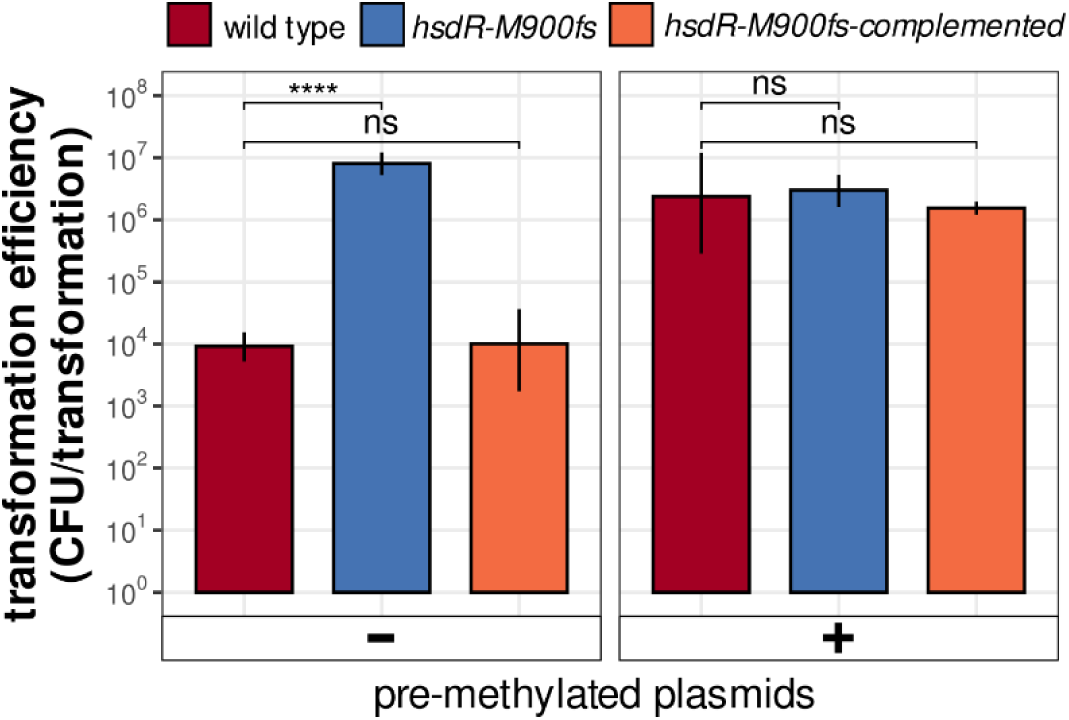
SSR-mediated frameshift in hsdR increases horizontal gene transfer in S. aureus. Transformation efficiency BPH2986 wild type, BPH2986-M900fs and complemented mutant. For transformation, plasmid pRAB11 (5 µg) was isolated from either E. coli strain DC10B (left) or IM08B strain to pre-methylate the plasmid and by-pass S. aureus T1-RM-system restriction of foreign DNA by hsdR (right)^100,96^. Bars represent mean transformation efficiency of three to four replicates and bars represent range of measurements (**** p≤0.0001, ns not significant, two-sided t-test).

### SSR-frameshift causes reversible small colony variant (SCV) phenotype switching

Small colony variants (SCVs) are an unstable *S. aureus* colony phenotype switch associated with persistent and relapsing *S. aureus* infections. SCVs are challenging to detect and treat ^17,43,44^. We identified a highly convergent SSR-frameshift in the *menF* gene, implicated in the menadione biosynthetic pathway (*menF-N186fs,* Supp. Table 1,3). As menadione auxotrophy has been reported in clinically isolated SCVs, we investigated if *menF-N186fs* could cause pathoadaptative SCVs ^45^. Macroscopic observation of the transposon mutant JE2*-menF::Tn* ^46^ confirmed that *menF* inactivation causes a SCV phenotype (Fig. 6a). Interestingly, the introduction of the most recurrent SSR-mediated *menF* frameshift in *S. aureus* BPH2986 led to a SCV phenotype (*menF-N186fs-(A)_6_-SCV*), that spontaneously reverted to Normal Colony Variant (NCV) upon cultivation (Fig. 6a and b). Whole genome sequencing of a NCV revealed the genotypic reversion of *menF* SSR to its wildtype form (*menF-N186fs-(A)_6→7_-NCV*). Menadione supplementation promoted normal colony growth of both the *menF::Tn* mutant and *menF-N186fs-(A)_6_-SCV*, confirming the consequence of *menF* inactivation on menadione auxotrophy (Fig. 6a). Phenotypic assessment showed that the *menF* SSR mutant (*menF-N186fs-(A)_6_-SCV*) was non haemolytic, whereas the subpopulation of spontaneous NCV revertant colonies (*menF-N186fs-(A)_6→7_-*NCV) were haemolytic like the wild type strain (Fig. 6b). The SSR-frameshift in *menF* also caused resistance to the aminoglycoside antibiotic gentamicin (MIC increase from 0.75 to 8 mg/mL, Fig. 6c). We next investigated the impact of *menF* mutants upon infection of epithelial cells. We continuously monitored HeLa cell viability post infection by measuring propidium iodide (PI) fluorescence ^11^. Both *JE2*-*menF::Tn* and *menF-N186fs-(A)_6_-SCV* mutants demonstrated significant loss of intracellular cytotoxicity compared to their cognate parental strains (p≤0.0001, two-sided Wilcoxon test; Fig. 6d). Infection of epithelial cells with the *menF* frameshift revertant (*menF-N186fs-(A)_6→7_-*NCV) restored intracellular cytotoxicity to the level of the wild type (Fig. 6d). These results demonstrate that SSR generates unstable, menadione auxotrophic SCVs known to cause persistent *S. aureus* infections ^45^. These results provide a mechanistic explanation of the historical observation of dynamic gentamicin-resistant SCV subpopulations arising in *S. aureus*, in absence of selective pressure ^47,48^. We also found that *menF-N186fs* subpopulation is non-haemolytic and has reduced intracellular cytotoxicity. These are phenotypes associated with staphylococcal persistence within host cells ^11^. In all, these data show a new role of SSRs in creating antibiotic-resistant intracellular reservoir of SCVs, contributing to chronic and recurrent infections.

**Figure 6.**
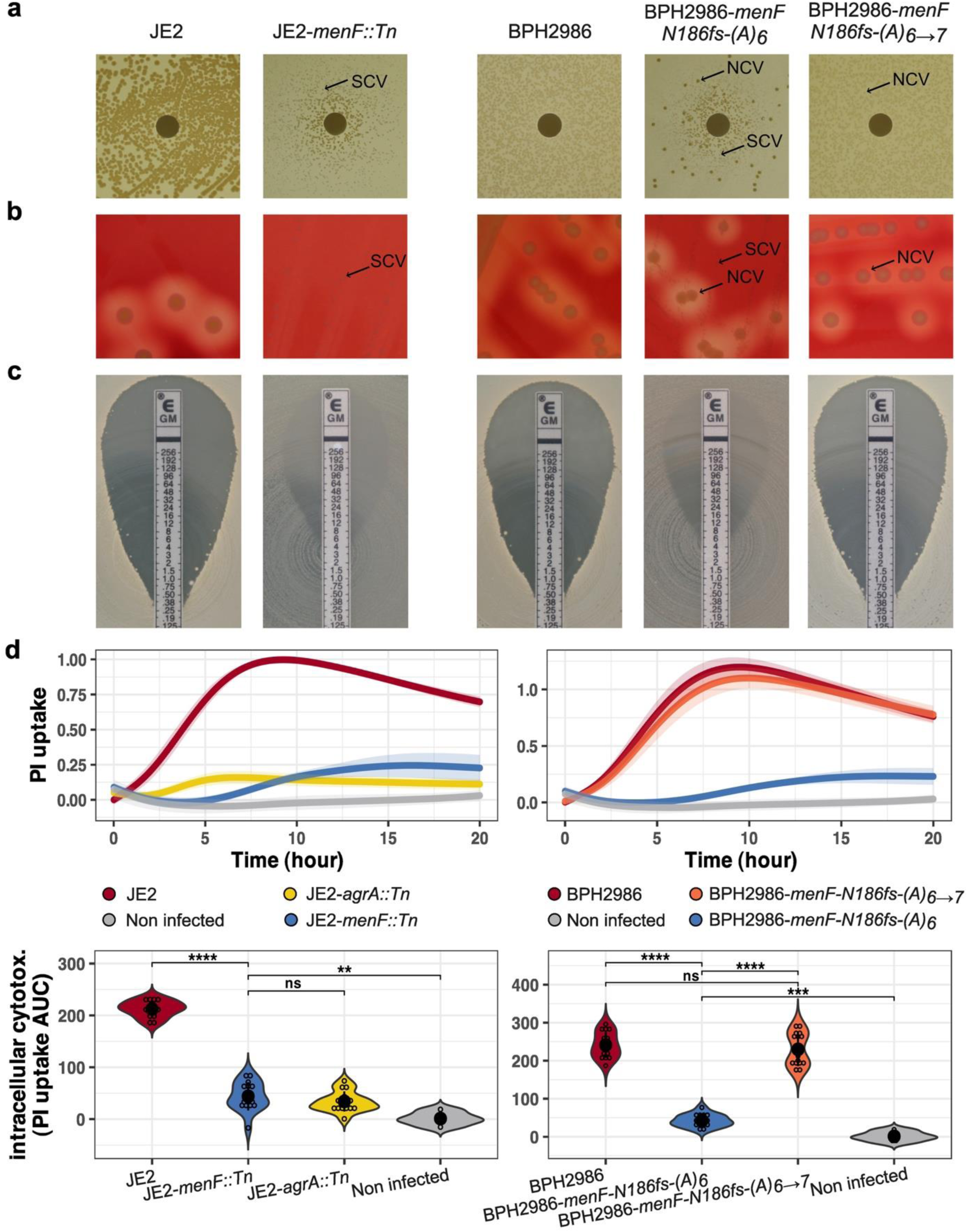
SSR mediated frameshift in menF causes unstable SCV phenotype. **a**. Menadione auxotrophy of menF mutants assessed by Menadione sodium bisulfite disc diffusion assay. **b**. Reduced haemolysis of menF mutants. **c**. Gentamycin resistance of menF mutants assessed by E-Test. **d**. Intracellular toxicity of S. aureus (InToxSa) assay ^11^ of menF mutants. Depicted PI uptake curves reflect Hela cells death post-infection with S. aureus. For each curve, the thick line represents the mean PI uptake signal across time of 14 replicates and shading around curves delineates the 95% confidence interval of the means. Violin plots represent the density distribution of the Area Under the Curve (AUC) of PI uptake of 14 replicates (** p≤0.01, *** p≤0.001, **** p≤0.0001, ns not significant, two-sided Wilcoxon test).

### Other SSR-mediated frameshifts have important impacts on adaptation

The experimental characterisation of three important SSR loci highlighted SSRs as key drivers of genome plasticity and adaptive evolution in *S. aureus*. These SSRs generate cell subpopulations with increased capacity to adapt (*mutL*, *hsdR*) or cause difficult to treat persistent infections (*menF* SCV). A review of the 171 genes impacted by recurrent SSR-frameshifts (Supp. Table 3) identified 17 genes for which important pathoadaptive phenotypes are predicted based on previous studies (Table 1). These SSR loci are notably predicted to generate heterogeneous population of *S. aureus* cells with: increased resistance to key antibiotics used to treat *S. aureus* infections (*tcaA*: glycopeptide ^49^, *lytH*: methicillin ^50^, *menF*: aminoglycoside); modulation of biofilm formation or exist as free-floating cells (*rbf* ^51^, *icaC* ^32^) and altered host-pathogen interactions (e.g. *capAC*: capsule ^52^, *eap/map*: adhesins ^53^, *essB*: type VII secretion system ^54^).

**Table 1.** SSR-mediated frameshift with high CMAS predicted to confer important adaptation. The references supporting the prediction of the impact of these SSRs can be found in Supp. Table 6. The full list of recurrent SSR-mediated frameshifts can be found in Supp. Table 1.

| Gene | CMAS | P | Product | Role | Predicted Pathoadaptation | SSR | Frameshift Mutation | Therapeutic target | Therapeutic type |
| --- | --- | --- | --- | --- | --- | --- | --- | --- | --- |
| agrA | 9.85 | 2.31E-04 | accessory gene regulator protein A | Quorum sensing<br>Transcription regulation | Intracellular persistence<br>Biofilm, Antimicrobial resistance, Virulence | (A)7 | I238fs | yes | - Vaccine<br>- Probiotic<br>- Bioactive compound |
| arlS | 5.62 | 2.88E-03 | sensor histidine kinase protein | Transcription regulation | Virulence<br>Host cell interaction | (T)7 | D265fs |  |  |
| atpA | 3.67 | 8.82E-03 | ATP synthase subunit alpha | ATP synthesis | Persisters cells generation | (T)7 | S497fs |  |  |
| capA | 7.10 | 1.19E-03 | capsular biosynthesis protein Cap5A | Capsule | Host cell interaction Immune evasion | (A)7 | N17fs | yes | - Vaccine<br>- Monoclonal Antibody |
| capC | 3.79 | 8.29E-03 | capsular biosynthesis protein Cap5C | Capsule | Host cell interaction Immune evasion | (A)7 | I237fs | yes | - Vaccine<br>- Monoclonal Antibody |
| crtP | 4.49 | 5.47E-03 | phytoene dehydrogenase | Staphyloxanthin production | Host cell interaction | (T)7 | N428fs | yes | - Bioactive compound |
| eap/map | 5.13 | 3.78E-03 | cell surface protein | Adhesin | Host cell interaction Immune evasion | (T)9 | N583fs | yes | - Vaccine |
| essB | 3.63 | 9.09E-03 | secretion system component EssB | Type VII secretion system | Interspecies competition<br>Host cell interaction | (A)7 | K444fs | yes | - Vaccine |
| fhuD2 | 4.32 | 6.11E-03 | hydroxamate siderophore binding lipoprotein | Iron transport | Immune evasion | (T)7 | N25fs | yes | - Vaccine |
| graX | 3.97 | 7.31E-03 | auxiliary protein GraX | Transcription regulation | Antimicrobial resistance<br>AMP resistance | (A)7 | V109fs |  |  |
| hssR | 5.96 | 2.40E-03 | DNA-binding heme response regulator | Transcription regulation | Host cell interaction | (A)7 | K203fs |  |  |
| icaC | 9.19 | 3.67E-04 | intercellular adhesion protein C | Polysaccharide intercellular adhesion biosynthesis | Biofilm formation<br>Immune evasion | (TTTA)3 | L287fs | yes | - Vaccine |
| lytH | 5.87 | 2.46E-03 | N-acetylmuramoyl-L-alanine amidase | N-acetylmuramoyl-L-alanine amidase | High level methicillin resistance | (T)7 | I4fs |  |  |
| rbf | 7.82 | 7.45E-04 | AraC family transcriptional regulator | Transcription regulation | Biofilm formation | (AT)5 | I51fs |  |  |
| ribA | 7.23 | 1.11E-03 | vitamin B2 biosynthesis protein | Vitamin B2 biosynthesis | Abolish MAIT cells recognition | (T)7 | I387fs |  |  |
| sarA | 5.64 | 2.83E-03 | accessory regulator A | Transcription regulation | Virulence<br>Host cell interaction | (T)6 | R84fs |  |  |
| tcaA | 4.38 | 5.87E-03 | teicoplanin resistance associated membrane protein | Cell wall | Antimicrobial resistance<br>Adaptation to bacteremia<br>AMP resistance | (T)7 | G390fs |  |  |

We also identified four major virulence-associated regulators (*agrA*, *sarA*, *arlS*, and *hssR*) as impacted by recurrent SSR frameshifts and four two-component regulatory systems (*agrA*, *arlS*, *hssR*, and *desK*). Two-component system regulators are used by bacteria to sense and respond to their environment ^55^. The SSR-mediated frameshifts affect regulators crucial for quorum sensing, toxin production, biofilm formation, exoprotein production, and *S. aureus* interaction with the host ^55^. Interestingly, the inactivation of virulence-associated regulators is frequently observed during invasive infections caused *by S. aureus* and other commensals ^12,27,56–58^. The recurrence of SSR frameshifts in these regulatory genes suggests that the ability to reversibly silence the typical bacterial response of subpopulations of cells favours survival in challenging conditions, such as invasive infections, where rapid adaptation to drastically different selective pressures is required.

Persister cells are a subset of bacterial cells that enter a dormant, non-growing state, allowing them to survive antibiotic treatment. Mutation of the *atpA* and *tcaA* genes, which encode the α subunit of the F1F0 ATP synthase and a membrane-associated protein, respectively, have been shown to promote persister cell formation ^59,60^. Interestingly, we identified a recurrent SSR-mediated frameshift in both *atpA* and *tcaA* genes, indicating that SSRs can also modulate the formation of persisters. Inactivation of TcaA, through the remodelling of the bacterial cell wall, is known to confer resistance to glycopeptide antibiotics and promote adaptation during blood infections by increasing resistance to serum and antimicrobial peptides. ^61^. Thus, by promoting co-adaptation to both antimicrobials and host defence during bacteraemia, the *tcaA* SSR-mediated frameshift would favour *S. aureus* survival in infected patients.

Mucosa-associated invariant T (MAIT) cells are thought to play an important, yet understudied role in the human immune system defence against *S. aureus* infections ^62,63^. We have identified a recurrent SSR that is predicted to create subpopulations of *S. aureus* capable of fully evading MAIT cells-mediated immunity. This SSR causes a frameshift mutation in the *ribA* gene, which encodes an enzyme of the riboflavin vitamin pathway which synthetises the intermediate molecule that permits recognition by MAIT cells (5-Amino-6-(5-phosphoribosylamino) uracil or 5-A-RU). This finding suggests that *S. aureus* has evolved mechanisms to escape MAIT cell-mediated immunity.

Despite promising results in clinical studies, unfortunately no vaccine has yet been approved for human use. An essential consideration in vaccine development lies in the choice of antigens that induce a robust immune response and are reliably expressed, enabling consistent recognition by the immune system. It has been postulated that the repeated failures in vaccine development may be partly due to the phenotypic switching of *S. aureus*. ^64^. Critically, our study identified recurrent SSR-mediated frameshifts in antigenic targets chosen in major vaccine efficacy clinical trials, such as: *S. aureus* capsule (*capA* and *capC* genes), type VII secretion system (*essB*) and iron transport lipoprotein (*fhuD2*) (Table 1) ^65–70^. In total, we identified eight *S. aureus* functions targeted by vaccines, monoclonal antibodies, bioactive compounds and probiotics in preclinical and clinical studies for anti-staphylococcal treatments (Table 1). Our findings reveal SSR-mediated frameshifts as a pervasive mechanism of antigen and therapeutic target variation in *S. aureus*, contributing to the challenges encountered during the development of effective therapeutics.

### *S. aureus* generates niche-specific SSR variants subpopulations in human derived samples

Our identification of a wide range of potentially adaptative SSR-mediated frameshifts prompted us to assess if heterogeneous populations of SSR variants were detectable in microbiome samples from human origin. We also postulated that due to the frequent fluctuations, many SSR loci might be heterogenous in clinical samples, with co-existing expanded and contracted populations. To investigate this, we searched for human-derived samples from which deep-population sequencing revealed a significant amount of *S. aureus* sequences. We analysed 2,244 metagenome samples from three distinct human origins: nasal swabs from healthy young donors, sputum from cystic fibrosis patients and blood culture from periprosthetic tissues positive for *S. aureus* ^71–73^. We identified 29 deep-metagenome samples from which *S. aureus* sequences were present with sufficient depth for the detection of *S. aureus* subpopulations (> 80% genome coverage and average sequencing depth > 50 reads, see method). Subpopulations of SSR variants were detected in 93% of these samples (27/29). A broad and diverse population of intragenic SSR variants was detected with up to 30 SSR variants per sample with allele frequencies ranging from 10 to 50% (Fig. 7a; Supp Table 7). Interestingly, the patterns of intragenic SSR variants differed between niches, with the number of SSR variants specific to sample origin outnumbering the number of SSR shared across niches (Fig. 7b). This analysis highlights the clinical relevance of SSRs, as subpopulation profiles most likely reflect niche-specific adaptations. Considering the extensive SSR-mediated population diversity identified, deep population sequencing appears necessary to fully capture its evolutionary dynamics within the host. Unlike single-isolate sequencing approaches, a population-level strategy has the potential to reveal adaptive phase-variation promoting colonisation and infection. Because SSR-mediated variation can influence *S. aureus* ability to evade immune responses, develop antibiotic resistance, and drive pathogenicity, mapping these trajectories could uncover new avenues for more effective disease management.

**Figure 7.**
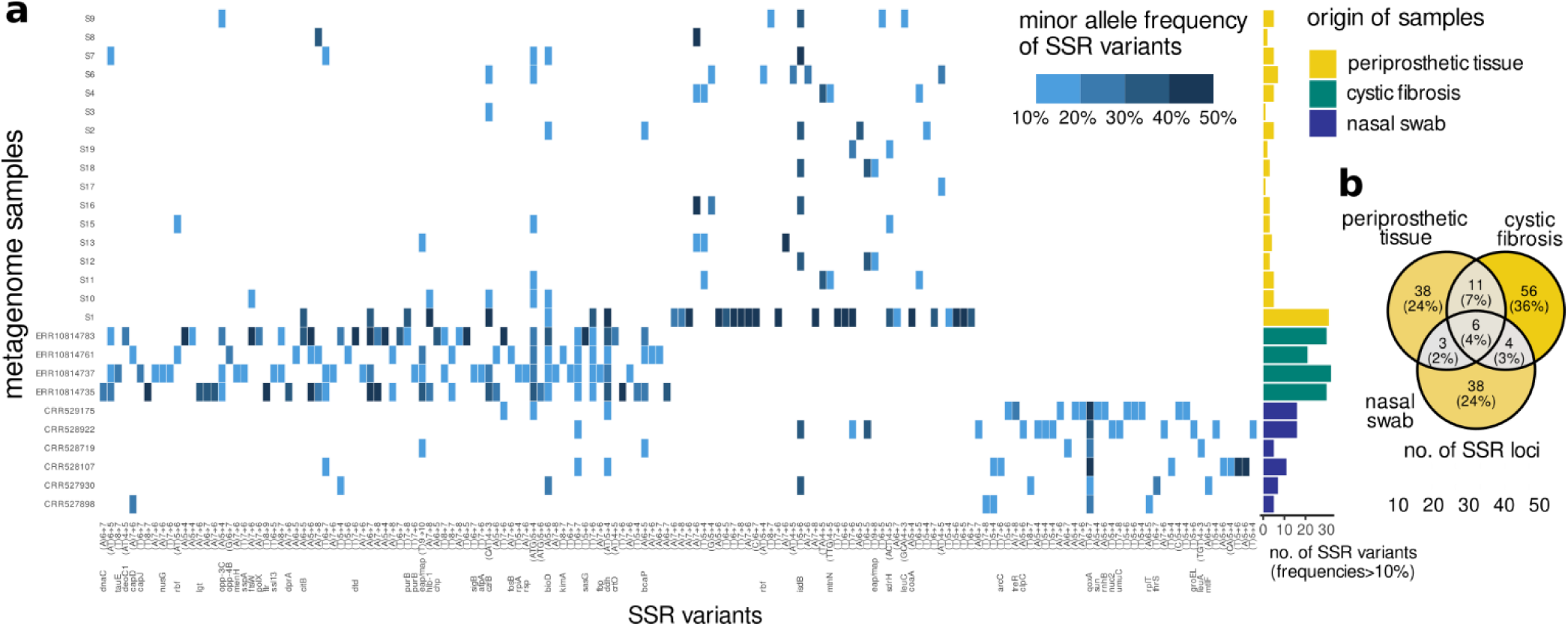
Heterogenous populations of SSR variants in human-derived deep-metagenome samples. **a**. Heatmap representing minor SSR alleles frequencies detected in 27 deep-population sequencing of human-derived samples. Minor SSR variants superior to 10% are represented with blue tiles (Supp. Table 7). Higher frequencies are indicated with darker blue. The bar chart on the right side of the heatmap indicates the number of minor SSR variants detected in each sample with colour indicating their origin. **b**. Venn diagram showing the number and percentage of SSR variants specific to or shared across the different sample origins. Metagenomes metadata and references are presented in Supp. Table 8.

## Discussion

*Staphylococcus aureus* is integral to the human microbiome but it is also the major cause of lethal bloodstream infections ^16^. In this study we developed an innovative, systematic *in silico* approach to identify adaptive *S. aureus* mutations. This screening method led to the discovery of the unexpectedly pervasiveness of SSRs as a major mechanism of *S. aureus* genetic variation.

We show that *S. aureus* has evolved hundreds of SSRs that in turn can generate extensive phenotypic heterogeneity. This bet-hedging strategy allows *S. aureus* populations to generate a spectrum of phenotypically diverse subpopulations, ensuring preparedness for rapid environmental fluctuations ^74^. When the environment and selective pressure on a population changes, subpopulations of pre-existing SSR variants better adapted to the new conditions can expand, eventually reaching a point of quasi-fixation within the population. This strategy enables rapid adaptation, conferring an exceptional ability for *S. aureus* to resist intense selective pressures and thrive in diverse, changing environments within the human body.

A striking discovery was that the *S. aureus* DNA repair gene *mutL* had one of the longest and most unstable SSRs. The loss-of-function frameshift mutation in this SSR significantly elevated *S. aureus* mutation rates. The *mutL* SSR signature was also detectable in a measurable fraction of *S. aureus* cells in otherwise clonal populations obtained from human colonisation and infection episodes. The SSR-mediated hypermutator phenotype is thus a gateway for increasing *S. aureus* genetic diversity and accelerating clinically significant phenotypes such as the development of antimicrobial resistance. With a long and unstable SSR in a DNA repair gene, *S. aureus* has evolved a strategy that maximises genetic diversity in a minor subset of the population while preserving genetic stability in the majority. This balance gives the bacterial population a large adaptive potential that can respond to rapid environmental changes, while minimising genome decay and loss of fitness.

Beyond hypermutation, this study shows that *S. aureus* employs SSRs to facilitate more specific adaptive strategies, including the capacity to remove the barrier to foreign DNA uptake and the formation of antibiotic-resistant subpopulation of small colony variants (SCVs), adapted for an intracellular lifestyle and causing persistent infection. Interestingly, prior to the genomic era, it has been reported that *S. aureus* populations continuously produce a small, dynamic subpopulation of gentamicin-resistant SCVs in the absence of selective pressure ^47,48^. The identification of *menF*-SSR fluctuation provides the mechanistic explanation for this phenomenon identified 20 years ago. SSR-mediated generation of SCV likely contributes to chronic and recurrent infections by generating an antibiotic-resistant intracellular reservoir that can persist in host tissues and revert to the parental phenotype.

Molecular mechanisms underlying persister cell formation, causing antimicrobial tolerance, are not well understood, although persisters in *S. aureus* have been associated with ATP depletion ^75,76^. Due to the high reversion rate of this phenotype, persister cell generation is thought to be inherited, not to driven by mutations ^77,76^. The SSRs variations reported here challenge that understanding, as the elevated mutation rate of SSRs permits the generation and reversion of growth-arrested subpopulations, characteristic of persister cells. This study also provides evidence that SSRs generate subpopulations favouring persisters (*atpA*, *tcaA*) ^59,60^.

An extensive SSR-mediated genetic diversification mechanism in *S. aureus* has several clinical implications. The potential for *S. aureus* to switch off or mutate key antigen genes and pathogenesis-related genes pose challenges for vaccine development and therapeutic interventions. Understanding these diversification mechanisms opens new avenues for research into more effective treatments. Additionally, deep-sequencing approaches that bypass traditional isolation cultivation methods can provide a more comprehensive understanding of bacterial pathoadaptation during infection. Future research should focus on exploring the potential of *S. aureus* SSR patterns in clinical isolates as prognostic markers of disease progression, with a vision to use that information to tailor treatments during infection.

While our study provides valuable insights into the adaptive mechanisms of *S. aureus*, it has some limitations to consider. Relying on metagenomic data for the identification of SSR variation *in vivo* from different clinical studies may introduce potential biases—such as differences in geographic origin, host demographics, sequencing platforms and protocols, and batch effects—that could influence observed SSR variant patterns. Nevertheless, the distinctive SSR subpopulation patterns identified across niches cannot be attributed solely to these biases and call for deeper investigation into how specific SSR variants correlate with clinical presentations and patient outcomes. Additional functional studies could also be conducted to assess the impact of the many highly recurrent SSR mutations detected through genomics but not assessed experimentally. *In vivo* studies in particular will be very interesting to undertake, to monitor changes in SSR-mediated diversity on adaptations under different infection conditions.

SSRs are predicted to predominantly confer loss-of-function (LoF) via frameshift mutations, which appears as limitation in their adaptive potential. Nevertheless, LoF mutations are becoming recognised as major contributors to rapid adaptation, and selection experiments in mutagenised bacteria demonstrate that substantial fitness gains can arise solely through LoFs ^78^. The “less-is-more hypothesis” posits that gene inactivation is not a byproduct of decay but represents a common adaptive strategy—particularly when populations encounter abrupt shifts in selective pressures and new environments ^79^. In *S. aureus*, LoF mutations appear as key drivers of patho-adaptation, exemplified by frequent inactivation of the major virulence regulator *agrA* in clinical isolates and the repeated observation of an overabundance of LoF mutations in invasive strains ^12,27^. By reprogramming regulatory networks or disabling metabolic pathways—such as those leading to small-colony variants or persisters—LoFs can confer substantial fitness advantages under drastic host and antibiotic pressures. However, classical LoF mutations are essentially irreversible, requiring rare back-mutations or compensatory changes and thus offering limited adaptive plasticity. In contrast, SSR-mediated LoFs generate reproducible on/off subpopulations for specific functions and revert readily when selection shifts—at rates up to 10^5^-fold higher than spontaneous point mutations. This dynamic ‘bet-hedging’ strategy is well suited to rapidly fluctuating environments but likely incurs costs in stable settings. Hence, while SSR-mediated LoFs can expand adaptive potential, its conservation within a given gene over the long term probably depends on how much selection on its function fluctuates.

In conclusion, this study reveals the importance of SSRs in understanding the adaptive strategies of *S. aureus*. The discovery of the extent of SSR-mediated genetic heterogeneity provides a new perspective on *S. aureus* adaptability, with profound implications for combating difficult-to-treat infections. Continued research into SSR patterns and their role in immune evasion and pathogenesis evolution will pave the way for rethinking therapeutic approaches and improving strategies to combat *S. aureus* infections.

## Methods

### Phylogenetic reconstruction

*S. aureus* genomes were downloaded from NCBI database (NCBI Bioproject list in Supp. Table 9). All sequence files were processed with *snippy* (v4.2; T. Seemann; https://github.com/tseemann/snippy) and mapped against fully assembled reference genome of *S. aureus* USA300 strain NRS384 (NCBI refseq assembly: GCF_008244705.1). High-confidence variants were called by removing aligned reads having mapping quality below 60 and requiring a minimum depth of 20 reads with at least 90% of reads supporting variants. To remove low-quality genome sequences, we kept only the genome sequences covering at least 80% of the reference genome. In total, we kept 7,099 *S. aureus* genomes for downstream analysis. To generate the core genome SNP phylogeny, whole genome alignment positions that were non monomorphic (at least one variant across all genomes) and with less than 2% gaps were retained using *snp-sites* (v2.3.2) and *trimAl* (v1.4.rev15), respectively ^80,81^. Maximum likelihood phylogenetic tree was constructed with RAxML (v8.2.9) ^82^ using the generalised time-reversible model with optimisation of substitution rates and gamma distribution to account for among site rate heterogeneity (using argument –m GTRCAX).

### Convergence analysis and estimation of the number of independent acquisitions of mutations

Using the phylogenetic reconstruction establishing the evolutionary history of the 7099 isolates from core genome SNP, a parsimony analysis was performed for each detected mutations to estimate the minimum number of independent, monophyletic acquisition events required to explain the observed distribution of each mutation across the tree tips (i.e., the isolates). An independent acquisition is defined as a distinct evolutionary event (monophyletic event) in which a specific mutation arises de novo in a lineage rather than being inherited from a common ancestor that already harbored that mutation. The R script *homoplastic_score* (https://github.com/rguerillot/Simple-sequence-repeats-are-major-drivers-of-adaptive-evolution-in-Staphylococcus-aureus) calculate the most parsimonious number of independent acquisition using the *phangorn* R package ^83^ as in ^84,85^.

### CMAS calculation

The CMAS value indicates how many standard deviations the count of independent mutation acquisitions at a specific locus deviates from the average number of all independent acquisitions for the affected gene. This standardisation of acquisition counts at the gene-level corrects for linkage disequilibrium due to recombination—where blocks of mutations co-transfer and would otherwise appear as convergent mutations—and prevents these linked acquisition events from inflating convergent adaptive signals. It also accounts for inherent variability in mutation rates across genes arising from gene length, divergence time, and functional constraints, ensuring that only loci with acquisition counts truly exceeding their expected variance are identified. This standardisation is calculated using the following formula:

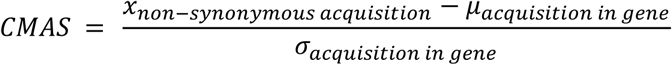

where: x non-synonymous acquisition is the observed count of independent non-synonymous mutation events at a specific locus; μ acquisition in gene is the mean number of independent mutation acquisition across the entire gene; and σ acquisition in gene is the standard deviation of mutation acquisition for that gene. Rather than assuming a specific model, the probability that a CMAS score equals or exceeds a given value is derived from the empirical distribution of scores across all genes of the *S. aureus* genome. Given that our dataset is very large and only a minimal fraction of mutations is expected to be adaptive, the empirical distribution robustly captures the true landscape of mutation acquisition frequencies. A nonparametric confidence interval ^86^ is constructed such that mutations with CMAS values exceeding the upper bound of the 99% prediction interval (a threshold of 3.464 in our dataset) are considered significant. This indicates that the mutation locus is acquired at a frequency far above neutral expectations and is likely under intense positive selection. For each mutation, we computed the probability that its CMAS score is equal to or exceeds its observed value. These empirical P values are provided in the “P” columns of Supplementary Tables 1, 2, 3, 6, and Table 1. CMAS was calculated after excluding position of the genome affected by frequent ambiguous mutation calls, frequent missing calls, or high rate of recombination. Highly recombinogenic region were identified by *gubbins* (v2.4.1) ^87^*. Gubbins* was run 8 times on random samples of 500 isolates using the bash script *sample_run_gubbins.sh* (https://github.com/rguerillot/Simple-sequence-repeats-are-major-drivers-of-adaptive-evolution-in-Staphylococcus-aureus). Results of *gubbins* runs were summed per position using *bedtools genomecov* (v2.30.0) ^88^ to obtain a recombination score per position. Positions with recombination scores <80 were retained for CMAS calculations. The recombination score threshold was determined by identifying the inflection point in the empirical cumulative distribution function of recombination scores. This threshold effectively excluded known mobile elements, such *S. aureus* prophages *Sa2* and *Sa3*, while retaining over 80% of the chromosome ^89^. The percentage of ambiguous and lack of alignment (gap) at every position was calculated from *snippy* full genome alignment output using the script *get_gap_ambiguous_snp_score.sh* (https://github.com/rguerillot/Simple-sequence-repeats-are-major-drivers-of-adaptive-evolution-in-Staphylococcus-aureus) that internally use *trimAl* (v1.4.rev15) ^81^ to calculate the percentage of missing alignment (gap position in the alignment) and ambiguous variant call (n position in the alignment). Positions with <20% gap and <10% ambiguous variant calls were kept for CMAS calculations.

### Annotation of genomes and SSR loci

*S. aureus* genomes were annotated using prokka ^90^ (v1.14.6) and RAST ^91^. Gene nomenclature and other annotations from aureowiki ^92^ were transposed on NRS384 reference genome using blastp reciprocal best hit ^93^ using USA300 strain FPR3757 (NCBI RefSeq assembly GCF_000013465.1). SSR loci were identified in *S. aureus* genome using kmer-ssr tool ^94^

### SSR mutants and reporter strains construction

SSR mutations were introduced by allelic exchange with the pIMAY-Z vector as described in ^11^. The upstream (amplicon 1) and downstream (amplicon 2) regions of SSR mutation were PCR amplified with primers with overlapping tails incorporating SSR mutations to be introduced (primers sequence in Supp. Table 10). After gel extraction of PCR products, splice by overlap extension PCR was performed to generate the insert containing the desired SSR mutations. Each insert was cloned into linearised pIMAY-Z vector by Seamless Ligation Cloning Extract (SLiCE) cloning ^95^. Each plasmid was separately transformed into *E. coli* strain IM08B ^96^ confirmed by colony PCR, then purified and transformed into strain BPH2986 (ST8 USA300) ^97^ or NRS384 by electroporation and the allelic exchange procedure was performed as previously described.

### Whole genome sequencing of constructed mutants and reporter strains

Reconstructed SSR mutants and SSR reporter strains (*ermC* amd *mutL-lacZ* fusion) were validated by whole-genome sequencing on an Illumina NextSeq 550 platform (Illumina) to confirm their genotype. SSR mutations and chromosomal reporter constructs were confirmed by de novo assembly using *shovill* (v3.14; T. Seemann; https://github.com/tseemann/shovill) and reads mapping onto the BPH2986 ^97^ or NRS384 reference genomes using *snippy* (v4.2; T. Seemann; https://github.com/tseemann/snippy). All strains constructed in this study are listed in Supp. Table 11 and whole genome sequence are available on NCBI website under BioProject number PRJNA1198637.

### Measurements of SSR expansions and contractions

To measure the rates of SSR expansions and contractions, overnight cultures of reporter strains (NRS384-*ermC*) were serially diluted to inoculate approximately 100 bacteria into 1 mL of Brain Heart Infusion Broth (BHIB) and grown for 25 generations. For this, strains were isolated from glycerol stocks onto Brain Heart Infusion Agar (BHIA). Independent colonies were inoculated in 10 mL of BHIB and incubated at 37°C overnight with agitation (200 rpm). The cultures were normalised to an OD600nm of 1.0 in BHIB and a 10-fold serial dilution to 10^-7^ was performed. In triplicate 1 mL of the 10^-7^ dilution and 1 mL of BHIB as a sterility control was aliquoted into a 24-well flat-bottom plate which was sealed with a plastic plate seal (ThermoFisher Scientific) and incubated for 24 h at 37°C with agitation (200 rpm). One hundred microliters of the 10^-7^ dilution was spread onto BHIA and incubated at 37°C overnight to calculate the colony-forming units per milliliter (CFU/mL) of the inoculum. After 24 h, to induce *ermC* expression, 10 μL of 0.01 mg/mL azithromycin (final concentration of 0.1 μg/ml) was aliquoted into each well and the plate was sealed and incubated for 2 h at 37°C with agitation (200 rpm). After 2 h, the bacterial suspension was serially diluted to 10^-7^. One hundred microlitres of the 10^-7^ dilution was spread onto BHIA to calculate the total CFU/mL of bacteria and 100 μL of the most appropriate dilution was plated onto BHIA containing 5 μg/ml azithromycin to calculate the subpopulation of azithromycin resistant bacteria in CFU/mL. All plates were incubated at 37°C for 24 h. Individual colonies were counted, and mutation frequencies were calculated by dividing CFUs on BHIA containing 5 μg/ml azithromycin plates by the total number of CFUs on non-selective BHIA plates. Mutation rates were calculated using the Drake formula ^98,99^.

### Assessment *mutL* SSR phase-variation by blue-white colonies screening

The reporter strain BPH2986-*mutL-lacZ-*N343fs-(A)_9_ was isolated from glycerol stock onto BHIA containing 1000 µg/mL X-Galactosidase (X-Gal) and incubated at 37°C. An independent colony was selected to inoculate BHIB and incubated at 37°C overnight with agitation (200 rpm). Five microlitres of overnight culture was sub-cultured in 10 mL of BHIB (Oxoid) and incubated at 37°C overnight with agitation (200 rpm). The culture was standardised at OD600nm of 1.0 and serially diluted 10^-4^ to 10^-5^ to obtain high cell density after plating. Using glass beads, a total volume of 800 µL of the diluted culture was spread evenly across square petri dishes of BHIA containing 1000 µg/mL X-Gal and incubated at 37°C overnight. Ten white colonies were selected for sequencing to further investigate the length of the SSR. The ten white colonies were colony purified on BHIA containing 1000 µg/mL X-Gal and genomic DNA (gDNA) was extracted from each colony using the GenElute Bacterial Genomic DNA Kit Protocol (Sigma Aldrich). The SSR region of *mutL* was amplified using primers Sanger_mutL_SSR_F and Sanger_mutL_SSR_R (Supp. Table 10) with 1 U Phusion High-Fidelity DNA Polymerase (ThermoFisher Scientific) with annealing at 55°C and extension at 72°C for 15 s for 25 cycles. The PCR products were purified using the FavorPrep Gel/PCR purification kit (Favorgen) and sequenced by Sanger sequencing method (Australian Genome Research Facility, Melbourne, VIC, Australia).

### Serial exposure to daptomycin and vancomycin

Strains were isolated from glycerol stocks onto BHIA. Independent colonies were inoculated into 500 μL Mueller-Hinton broth (MHB) for vancomycin testing or MHB with 50 μg/mL CaCl_2_ (cation adjusted) for daptomycin testing due to daptomycin calcium dependency. This suspension was diluted to 0.5 McFarland (McF). The 0.5 McF suspension was diluted 1:100 in MHB or cation adjusted MHB in a total volume of 1 mL. In a 96-well round bottom plate, 100 μL of MHB was aliquoted into column two to six and 200 μL of MHB was added to column seven as a sterility control. Two hundred microliters of MHB containing double the highest concentration of antibiotic to be tested was aliquoted into column 1, and a two-fold serial dilution was performed. Each well was inoculated with 100 μL of the 1:100 dilution of bacterial suspension, and the plate was incubated at 37°C for 18 to 22 h without agitation. After incubation the minimum inhibitory concentration (MIC) was determined. The suspension in the well with concentration just below the MIC well (0.5xMIC) was diluted to 0.5 McF then diluted 1:100 in MHB. A new serial exposure was set up in a 96-well plate as specified previously. New antibiotic exposure plates were incubated at 37°C for 18 to 22 h without agitation. Strains were serially exposed to vancomycin and daptomycin five times over five days in three independent experiments.

### RM-seq amplicon deep-sequencing of SSR loci

Genomic DNA was extracted from 1 mL of overnight BHI culture using the DNeasy Blood and Tissue Kit (QIAGEN) according to the manufacturer’s guidelines for lysis of gram-positive bacteria with the following modifications: 5 μL of 10 mg/mL RNase A (MP Biomedicals) and 1 μL 10 mg/mL lysostaphin (Ambi), instead of lysozyme, were added to the lysis buffer for each reaction. The RM-seq library was prepared as described in ^36^ with the following modifications. To introduce the random 16 bp barcodes 10 cycles of linear PCR was performed, and the PCR mix was as specified but contained 50 nM (ermC_rmseq_F or mutL_rmseq_F, Supp. Table 10) and 5 ng/μL of gDNA. For the nested exponential PCR, 3.25 μL of primer mix was added which contained 2 μL of 100 nM RM-seq reverse primer (ermC_rmseq_R or mutL_rmseq_R), 0.625 μL forward and 0.625 μL reverse 10 μM Nextera XT Index Kit primers (Illumina). The amplicons were purified using 0.9xPCR volume of Agencourt® AMpure® XP Magnetic Beads (Beckman Coulter) according to the manufacturer’s guidelines. The amplicons were normalised to 4 nM according to concentration and expected amplicon size, before being pooled. The sequencing library was diluted to 15 pM and the Reagent Kit v3 (Illumina) was used to prepare the sequencing libraries according to the manufacturer’s guidelines with the addition of 10% PhiX library. Sequencing using 2 by 150 bp paired-end chemistry was performed on the Illumina NextSeq 550 platform. Amplicons were sequenced and copy corrected using the rmseq analysis pipeline ^36^ and then SSR of different sizes were counted using the *rmseq_SSR_count.sh* script (https://github.com/rguerillot/Simple-sequence-repeats-are-major-drivers-of-adaptive-evolution-in-Staphylococcus-aureus).

### Within-host evolution analysis

We investigated a collection of 2,189 publicly available, sequentially collected *S. aureus* genomes from 190 episodes of airway infection in cystic fibrosis and 436 episodes of nose colonisation. Reads and metadata were retrieved from NCBI bioprojects listed in Supp. Table 9, when available. Reads were assembled using *shovill* and classified using *kraken*. Reads with a coverage below 20, less than 50% *S. aureus* reads and an assembly size greater than 3.5 Mb were excluded. Within-host evolution genomic analysis was performed as described in ^12^. Briefly, within-host variants in same-patient genomes were identified using *snippy* using the draft assembly of a randomly selected strain as internal reference. We excluded variants from internal reference reads and those in positions where the internal reference had a coverage below 10. To harmonise annotation, we clustered all internal reference coding regions using *cd-hit* and identified a homolog on reference genome *S. aureus* FPR3757 using *blastp*. The significance of within-host mutation enrichment at individual *mutL* amino acid positions was tested by Poisson regression as described in ^12^. To investigate whether isolates with a *mutL*-N343-frameshift had an accumulation of frameshifts, we fitted a multiple linear regression where the dependent variable was the total number of within-host acquired frameshifts, and the predictors were the *mutL* status of the isolate and the total number of mutations per isolate. To exclude episodes where reinfection could potentially be caused by a different *S. aureus* isolate, genomes with more than 100 within-host mutations were excluded from the regression (Supp. Table 5; total 1,355 isolates, cystic fibrosis [190 episodes, 592 genomes], nasal colonisation [436 episodes, 763 genomes]). We used *IQ-tree* (v2.0.3) to reconstruct the phylogeny of 7 infection episodes with de novo (within-host) emergence of the *mutL*-N343-frameshift and at least 4 sequences available per episode. We used *mash* to identify the closest genome obtained from another episode included in the same study and used it as an outgroup to root the tree. Trees were visualized using *ggtree*.

### Measurement of transformation efficiency

*S. aureus* cells were electroporated as described in ^100^. Five µg of the plasmid pRAB11 extracted from *E. coli* strain DC10B ^100^ or IM08B (for T1-RMS pre-methylation) ^96^ was used for electroporation of *S. aureus* BPH2986 and BPH2986-*hsdR-M900fs* strains in triplicate. After electroporation 1 mL of BHIB supplemented with 500 mM sucrose was immediately added to the cells which were incubated a 37°C for 1 h to allow cell recovery and expression of the chloramphenicol resistance gene before being plated onto BHIA containing 10 μg/mL chloramphenicol.

### Hela cells infection

Hela cells infection with *S. aureus* was performed using our *InToxSa* method as described previously^11^. All strains were assessed using 14 replicates over two different plates assessed during different days.

### Phenotypic assessment of SCVs

For the haemolysis assay, strains were isolated from glycerol stocks onto sheep blood agar (SBA) and incubated at 37°C overnight. For each strain an independent colony was subcultured onto SBA and incubated at 37°C overnight and the haemolysis pattern was observed. For Menadione Sodium Bisulfite (MSB) Disc Diffusion Assay, strains were isolated from glycerol stocks onto horse blood agar (HBA) or BHIA (BD Bacto). An independent colony was inoculated into 10 mL BHIB (BD Bacto) and incubated at 37°C overnight with agitation (200 rpm). The culture was standardised at OD600nm of 1.0 and serially diluted to 10^-4^. One hundred microlitres of the 10^-4^ dilution was spread plate onto BHIA (BD Bacto). A disc supplemented with 15 µL of 100 mg/L menadione sodium bisulfite (MSB, Sigma-Aldrich) was added to each plate and the plates were incubated at 37°C overnight. For Gentamicin Etest, strains were isolated from glycerol stocks onto SBA and incubated at 37°C overnight. For each strain an independent colony was subcultured onto HBA and incubated at 37°C overnight. A 0.5 McFarland Standard was setup by resuspending independent colonies in saline. BHIA (BD Bacto) was inoculated with the 0.5 McF standard, dried and gentamicin Etest strips (Biomérieux) were applied. The plates were incubated at 37°C for 24 h.

### Identification of SSR variants subpopulations in human derived samples

Sequencing reads from the six BioProjects listed in Supp. Table 9 were processed with the *bash* script *hetero_SSR_call.sh* available at (https://github.com/rguerillot/Simple-sequence-repeats-are-major-drivers-of-adaptive-evolution-in-Staphylococcus-aureus). Briefly, all fastq read files were mapped using *bwa* onto the *S. aureus* NRS384 reference genome. Sequencing depth and coverage were calculated *wi*th *samtools* (v1.10). Only samples with >80% *S. aureus* genome coverage and a mean sequencing depth of 20 were kept for further analysis. Then, *lofreq* ^101^ (v2.1.2), a pipeline specifically designed for the accurate calling of low frequency variants was used. SSR insertion deletion calls were further filtered to retain only minor variant call >10%.

## End notes

## Acknowledgments

We thank Tania Wong Fok Lung for helpful discussions.

## Author contributions

Romain Guérillot conceived the study, analysed the data, supervised the experimental work, generated the figures, and fully wrote the original draft of the manuscript. Ashleigh Hayes performed SSR mutants, SSR reporters, and RM-seq experiments and analyses, and assisted with the writing of the manuscript and the creation of figures. Abderrahman Hachani conducted the infection experiments, wrote the manuscript, and contributed to the study design and interpretation. Stefano G. Giulieri performed the within-host evolution analysis, created figures, wrote the manuscript, and contributed to the study design and interpretation. Liam Donovan contributed to SSR reporter experimental work. Louise M. Judd performed whole-genome sequencing. Liam K. Sharkey advised and contributed to the interpretation of the experimental work and analyses. Anders Gonçalves da Silva and Torsten Seemann advised and contributed to the bioinformatics analyses. Timothy P. Stinear and Benjamin P. Howden supervised the study design and interpretations, wrote the manuscript, and acquired funding for the study.

## Ethics declarations

### Availability of biological materials

Biological materials can be obtained by contacting the corresponding authors.

## Additional Information

Supplementary Information is available for this paper.

## Data availability

All data generated or analysed during this study are included in the manuscript and supplementary tables and figures. All publicly available genome and metagenome sequences used for the convergence evolution, within-host evolution and SSR heterogeneity analyses are available from the BioProjects listed in Supp. Table 9. Genome sequence generated for this study are available under NCBI BioProject number PRJNA1198637. The scripts used for analyses are available on the GitHub website at https://github.com/rguerillot/Simple-sequence-repeats-are-major-drivers-of-adaptive-evolution-in-Staphylococcus-aureus.

## Extended data figures and tables

**Supp. Figure 1.**
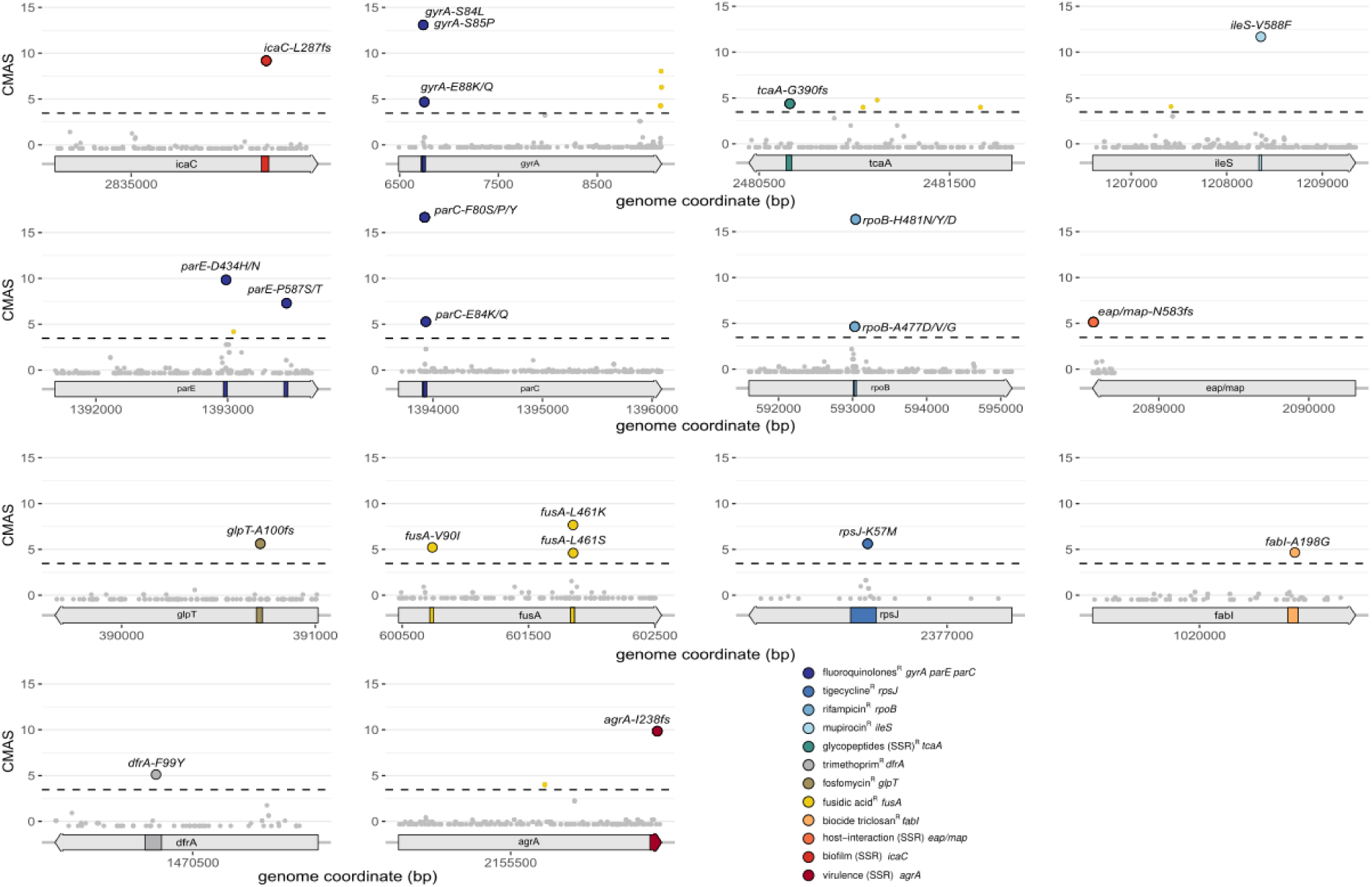
Gene-wide CMAS plots of 14 genes known to confer adaptive phenotypes. Mutations with significantly high CMAS (>99% prediction interval, dashed line) are represented by yellow dots (non-synonymous mutations). Mutations known to confer adaptive phenotypes are indicated with coloured circles with corresponding phenotypes indicated in the legend.

**Supp. Figure 2.**
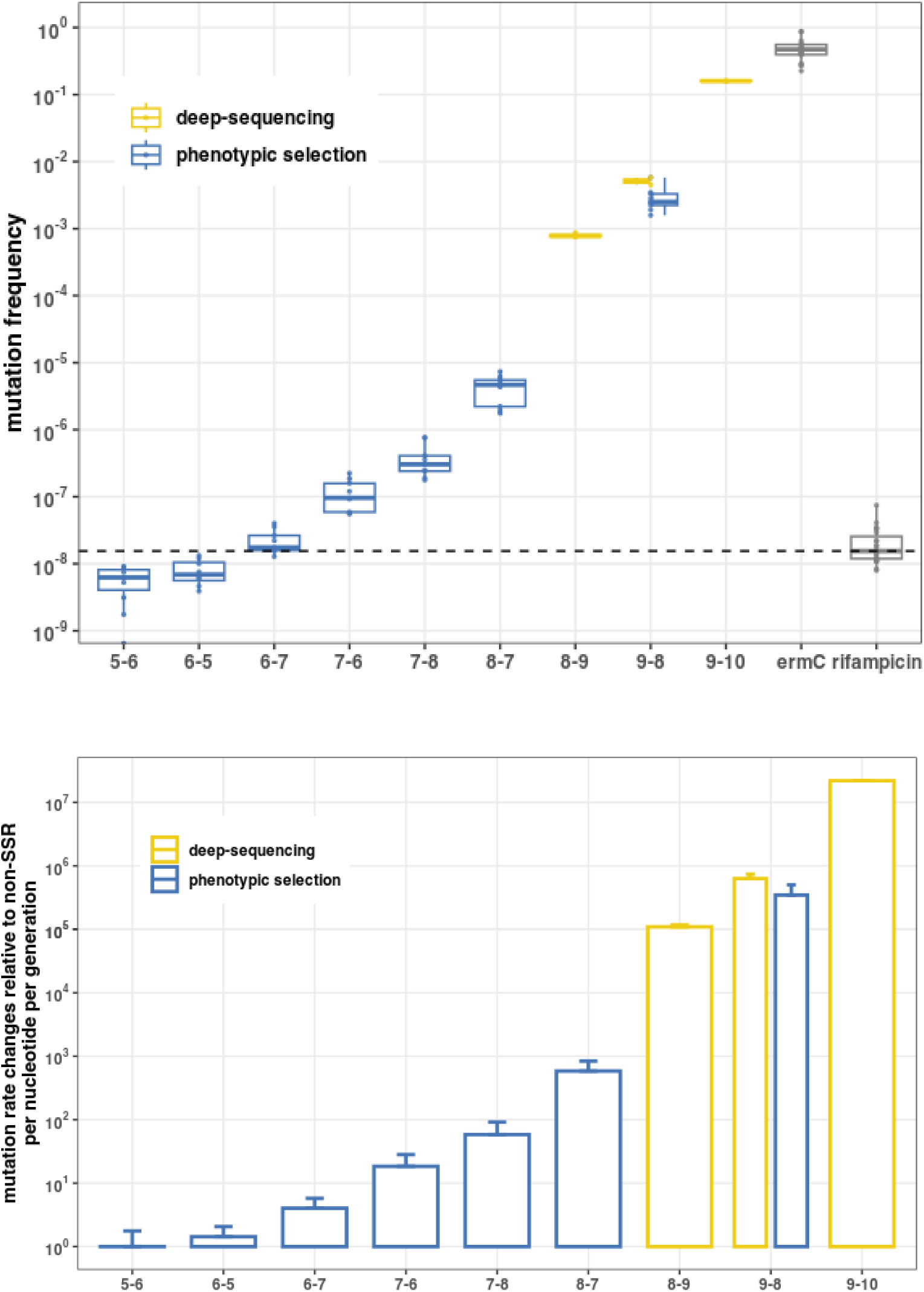
**a** Mutation frequency of the 7 SSR expansions (exp.) and contractions (cont.) assessed by azithromycin resistance selections (blue) and amplicon deep-sequencing (yellow). The ermC^wt^ allele is used as control and frequency of selection or rifampicin resistance is shown for comparison. **b.** Fold changes of mutation rate of SSR expansions and contractions relative to basal (non-SSR) mutation rate of S. aureus. Bars represent means and error bars represent standard deviations of change in mutation rate per nucleotide and per generation.

***Supp. Table 1. Convergent non-synonymous mutations.*** *All intragenic and non-synonymous mutations with significant CMAS.*

***Supp. Table 2. Previously described adaptive mutations rediscovered using CMAS.*** *This table list the 45 mutations in 14 genes known to confer adaptive phenotypes: resistance to 8 different antibiotic families, resistance to the biocide triclosan, virulence, modulation of biofilm formation and host-interaction.*

***Supp. Table 3. Convergent SSR mutations.*** *Intragenic and non-synonymous mutations affecting SSR with significant CMAS.*

***Supp. Table 4. Mutations associated with vancomycin and daptomycin resistance.*** *Mutations identified by whole genome sequencing of isolates obtained after the final exposure to sub-inhibitory concentrations of antibiotics.*

***Supp. Table 5. Within-host acquisition of the SSR mutation mutL-N343fs.*** *Sheet 1. Isolates with the mutL-N343fs mutation acquired during episodes of nasal colonisations or infections. Sheet 2. Total number of mutations and number of frameshifts mutations acquired within-host.*

***Supp. Table 6. References supporting the predictions of the adaptive impact of SSR-mediated frameshifts.*** *All SSR-mediated frameshifts with a significant CMAS.*

***Supp. Table 7. List of SSR variants in human-derived deep-metagenome samples.*** *Minor SSR alleles frequencies detected in 27 deep-population sequencing of human-derived samples. All SSR variants presented in the heatmap (Fig.7.a.) are listed with their annotation and their coordinate in S. aureus NRS384 genome.*

***Supp. Table 8. Metagenome data analysed for the detection of sub-populations of SSR variants.*** *The NCBI accession numbers, references and associated metadata of the metagenome samples with >80% S. aureus genome coverage and a mean S. aureus sequencing depth >20 reads.*

***Supp. Table 9. Publicly available genomic data analysed.*** *All project accession and project titles are listed. Sheet 1. Large scale convergence analysis of S. aureus genomes. Sheet 2 Within-host evolution of S. aureus. Sheet 3. SSR-driven heterogeneity in human-derived metagenome.*

***Supp. Table 10. Primers used in this study***

***Supp. Table 11. S. aureus strains and mutants reconstructed in this study***

