## Supplementary figures and images for "Simple sequence repeats power *Staphylococcus aureus* adaptation"

### Supp. Figure 1

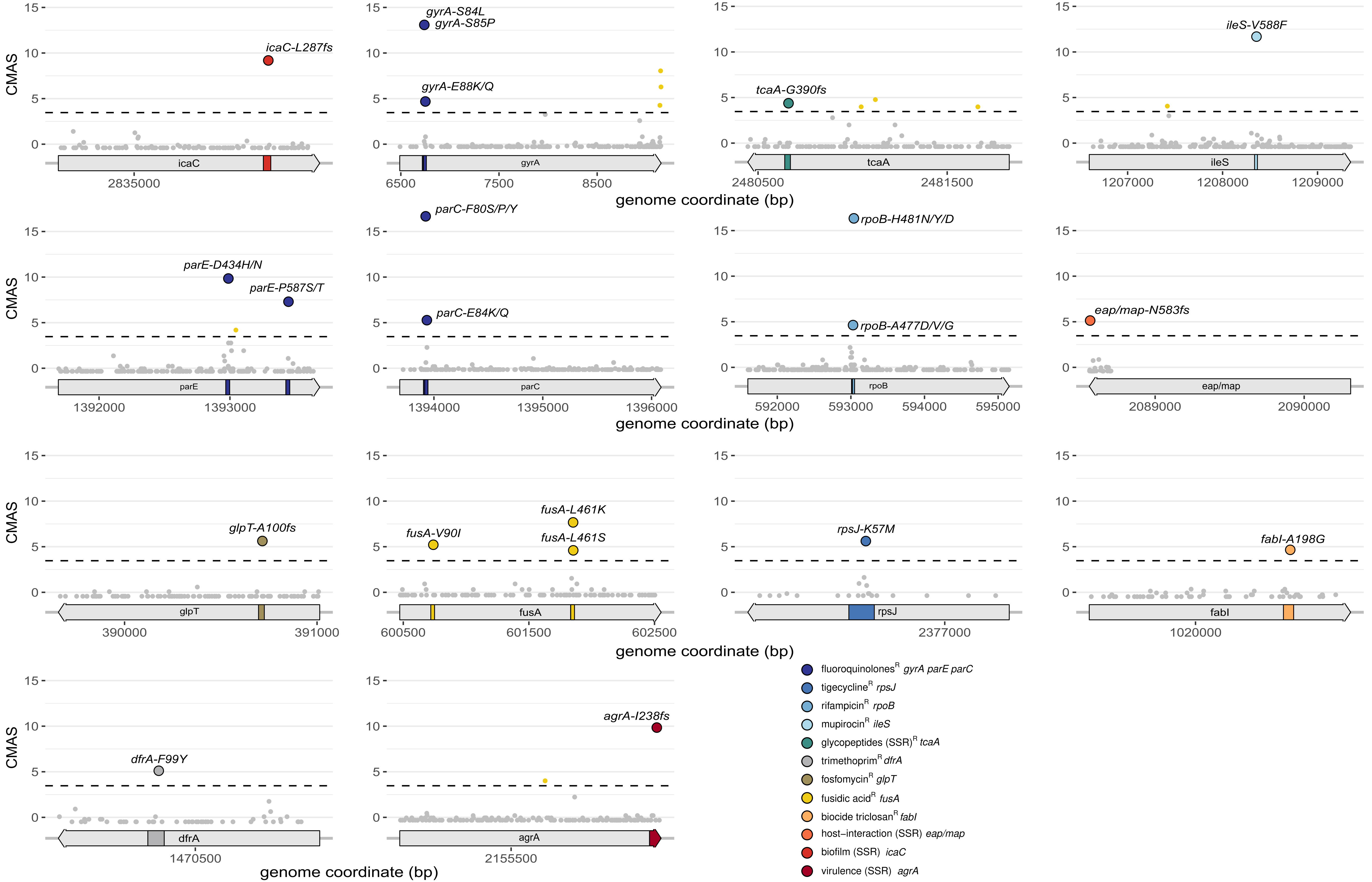

### Supp. Figure 2

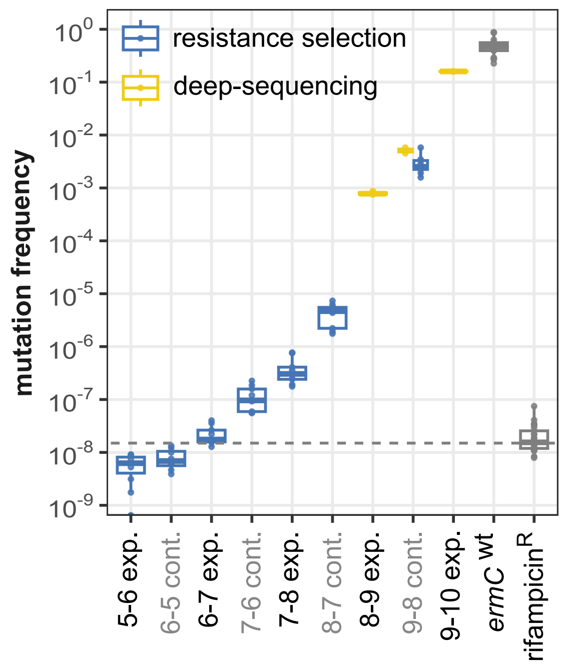
